# Experimentally induced active and quiet sleep engage non-overlapping transcriptional programs in *Drosophila*

**DOI:** 10.1101/2023.04.03.535331

**Authors:** Niki Anthoney, Lucy A.L. Tainton-Heap, Hang Luong, Eleni Notaras, Amber B. Kewin, Qiongyi Zhao, Trent Perry, Philip Batterham, Paul J. Shaw, Bruno van Swinderen

## Abstract

Sleep in mammals can be broadly classified into two different physiological categories: rapid eye movement (REM) sleep and slow wave sleep (SWS), and accordingly REM and SWS are thought to achieve a different set of functions. The fruit fly *Drosophila melanogaster* is increasingly being used as a model to understand sleep functions, although it remains unclear if the fly brain also engages in different kinds of sleep as well. Here, we compare two commonly used approaches for studying sleep experimentally in *Drosophila*: optogenetic activation of sleep-promoting neurons and provision of a sleep-promoting drug, Gaboxadol. We find that these different sleep-induction methods have similar effects on increasing sleep duration, but divergent effects on brain activity. Transcriptomic analysis reveals that drug-induced deep sleep (‘quiet’ sleep) mostly downregulates metabolism genes, whereas optogenetic ‘active’ sleep upregulates a wide range of genes relevant to normal waking functions. This suggests that optogenetics and pharmacological induction of sleep in *Drosophila* promote different features of sleep, which engage different sets of genes to achieve their respective functions.

## Introduction

There is increasing evidence that sleep is a complex phenomenon in most animals, comprising of distinct stages that are characterized by dramatically different physiological processes and brain activity signatures [1, 2]. This suggests that different sleep stages, such as rapid eye movement (REM) and slow-wave sleep (SWS) in humans and other mammals [3] are accomplishing distinct functions that are nevertheless collectively important for adaptive behavior and survival [4]. While REM and SWS appear to be restricted to a subset of vertebrates (e.g., mammals, birds, and possibly some reptiles [5-7] a broader range of animals, including invertebrates, demonstrate evidence of ‘active’ versus ‘quiet’ sleep [1, 2, 8]. Whether active and quiet sleep represent evolutionary antecedents of REM and SWS, respectively, remains speculative [1, 2]. However, during active sleep, although animals are less responsive, brain recordings reveal a level of neural activity that is similar to wakefulness, in contrast to quiet sleep, which is characterized by significantly decreased neural activity in invertebrates [9, 10] as well as certain fish [11], mollusks [12], and reptiles [6].

Although it is likely that even insects such as fruit flies and honeybees sleep in distinct stages [13, 14], sleep studies using the genetic model *Drosophila melanogaster* still mostly measure sleep as a single phenomenon, defined by 5 minutes (or more) of inactivity [15, 16]. As sleep studies increasingly employ *Drosophila* to investigate molecular and cellular processes underpinning potential sleep functions, this simplified approach to measuring sleep in flies carries the risk of overlooking different functions accomplished by distinct kinds of sleep. Sleep physiology and functions are increasingly being addressed in the fly model by imposing experimentally controlled sleep regimes, either pharmacologically or via transient control of sleep-promoting circuits by using opto- or thermogenetic tools [17]. Yet, there is little knowledge available on whether these different approaches are producing qualitatively similar sleep. For example, sleep can be induced genetically in flies by activating sleep-promoting neurons in the central complex (CX) – a part of the insect brain that has been found to be involved in multimodal sensory processing [18]. In particular, the dorsal fan-shaped body (dFB) of the CX has been proposed to serve as a discharge circuit for the insect’s sleep homeostat, whereby increased sleep pressure (e.g., due to extended wakefulness) alters the physiological properties of dFB neurons causing them to fire more readily and thereby promote decreased behavioral responsiveness [19] and thus sleep [20-22]. Crucially, activation of sleep-promoting circuits including the dFB have been shown to be sleep-restorative [10, 23], but confusingly, brain recordings during optogenetically-induced sleep, via electrophysiology as well as whole-brain calcium imaging techniques, reveal wake-like levels of brain activity [9, 10]. This suggests that some approaches to optogenetically-induced sleep might be promoting a form of sleep akin to the ‘active’ sleep stage detected during spontaneous sleep [9, 10].

An alternate way to induce sleep in *Drosophila* is by feeding flies drugs that have been designed to treat insomnia in humans, such as the GABA-agonist 4,5,6,7- tetrahyrdoisoxazolopyridin-3-ol, (THIP), also known as Gaboxadol [24]. Several studies have shown that THIP-induced sleep in flies is also restorative and achieves key functions ranging from memory consolidation to cellular repair and waste clearance [23, 25-27]. This pharmacological approach centered on GABA function has a solid foundation based on better-understood sleep processes: in mammals, many sleep-inducing drugs also target GABA receptors, and this class of drugs tends to promote SWS [28]. In contrast, there are no obvious drugs that promote REM sleep, although local infusion of cholinergic agonists (e.g., carbachol) to the brainstem has been shown to induce REM-like states in cats [29].

In this study, we compare THIP-induced sleep with optogenetically-induced sleep in *Drosophila*, using behavior, brain activity, and transcriptomics. To ensure the validity of our comparisons, we performed all of our experiments in the same genetic background, employing a canonical Gal4 strain that expresses a transgenic cation channel in a sleep-promoting circuit: R23E10-Gal4 > UAS-Chrimson [30, 31]. When these flies are fed all-trans-retinal (ATR) and then exposed to red light, they are put to sleep optogenetically. When these flies are instead fed THIP, they are put to sleep pharmacologically. By using the same genetic background, we were thus able to contrast the effects of either kind of sleep at the level of behavior, brain activity, and gene expression (**Figure 1**). We questioned how similar either form of induced sleep was.

**Figure 1.**
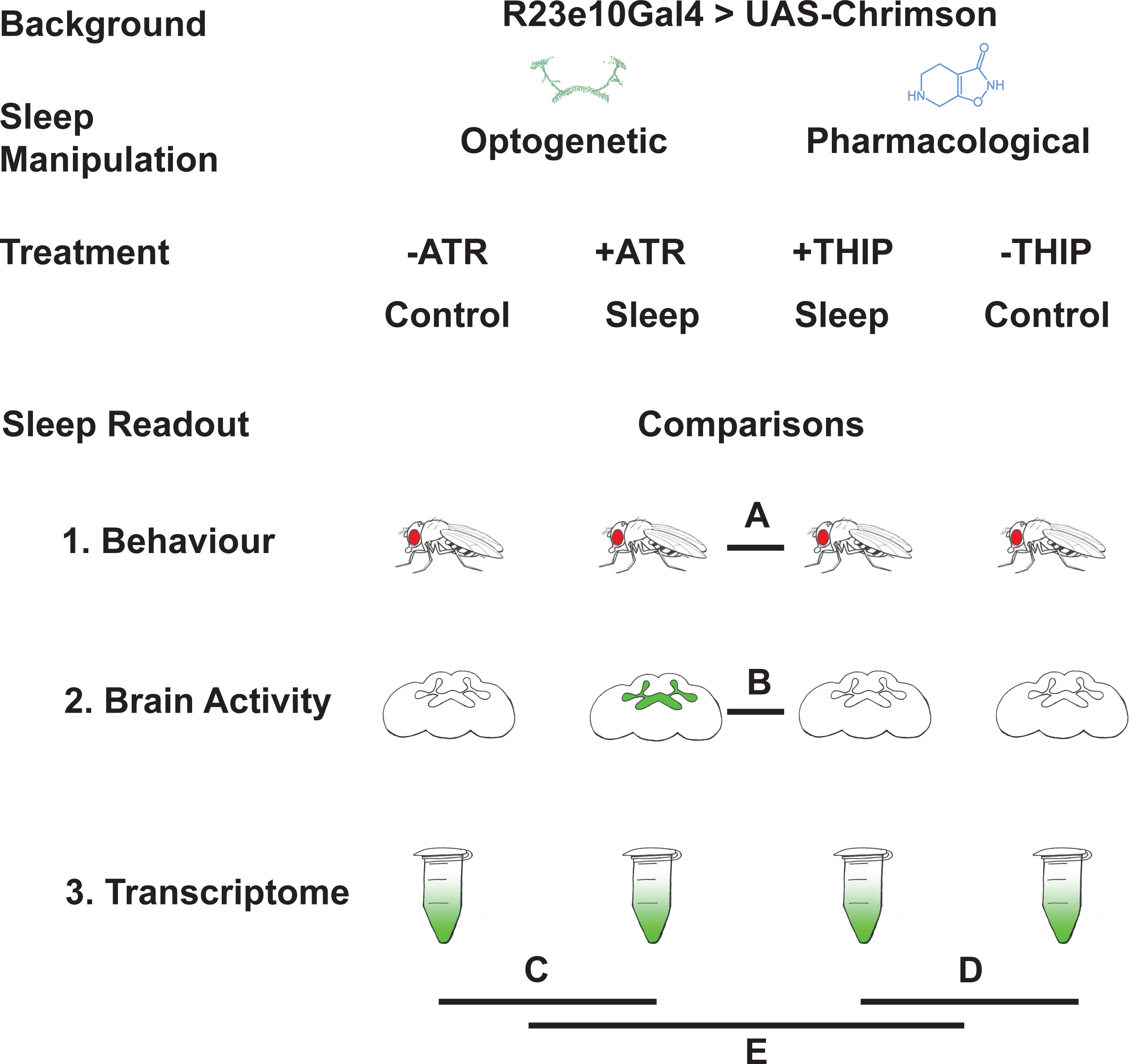
Study rationale and design. The same genetic background strain (R23E10;UAS-Chrimson) was used for optogenetic or pharmacologically-induced sleep. Flies were fed either all-trans retinal (ATR) or 4,5,6,7-tetrahyrdoisoxazolopyridin-3-ol (THIP) to promote either kind of sleep, which was assessed in three different ways: behavioral analysis, whole brain imaging, and gene expression changes. The comparisons made for each level of analysis are labelled A-E.

## Results

### Prolonged optogenetic and THIP-induced sleep have near identical effects on sleep duration

We first compared pharmacological and optogenetic sleep (**Figure 1**) by using the traditional behavioral metrics employed by most *Drosophila* sleep researchers: >5 minutes inactivity for flies confined in small glass tubes over multiple days and nights [15, 16]. We found that optogenetic- and THIP-induced sleep yielded almost identical effects on sleep duration, with both significantly increasing total sleep duration for both the day and night, when compared to controls (**Figure 2A-D; Supplementary File 1**). An increase in total sleep duration can be due to either an increase in the number of sleep bouts that are occurring (reflective of more fragmented sleep), or an increase in the average duration of individual sleep bouts, which indicates a more consolidated sleep structure [32-34]. To investigate whether both sleep induction methods also had similar effects on sleep architecture, we plotted bout number as a function of bout duration for optogenetic and THIP-induced sleep, for the day and night [35]. We found that both optogenetic activation and THIP provision produce a similar increase in sleep consolidation during the day (**Figure 2E, F; Figure 3**). During the night, induced sleep effects were also similar, although less clearly different to the spontaneous sleep seen in control flies (**Figure 2G, H; Figure 3**). Interestingly, red light exposure decreased average night bout duration in non-ATR control flies (**Figure 3A-D**), suggesting a light-induced artefact at night. For THIP, we observed an increase in both bout number and duration during the day, and an increase in bout duration during the night (**Figure 3E,F**). Taken together, these results show that prolonged optogenetic activation and THIP provision have similar behavioral effects on induced sleep duration in *Drosophila*, although some differences in night bout architecture were noted (**Figure 3**). Without any further investigations, this might suggest that both sleep induction methods represent similar underlying processes that uniformly increase sleep in flies.

**Figure 2.**
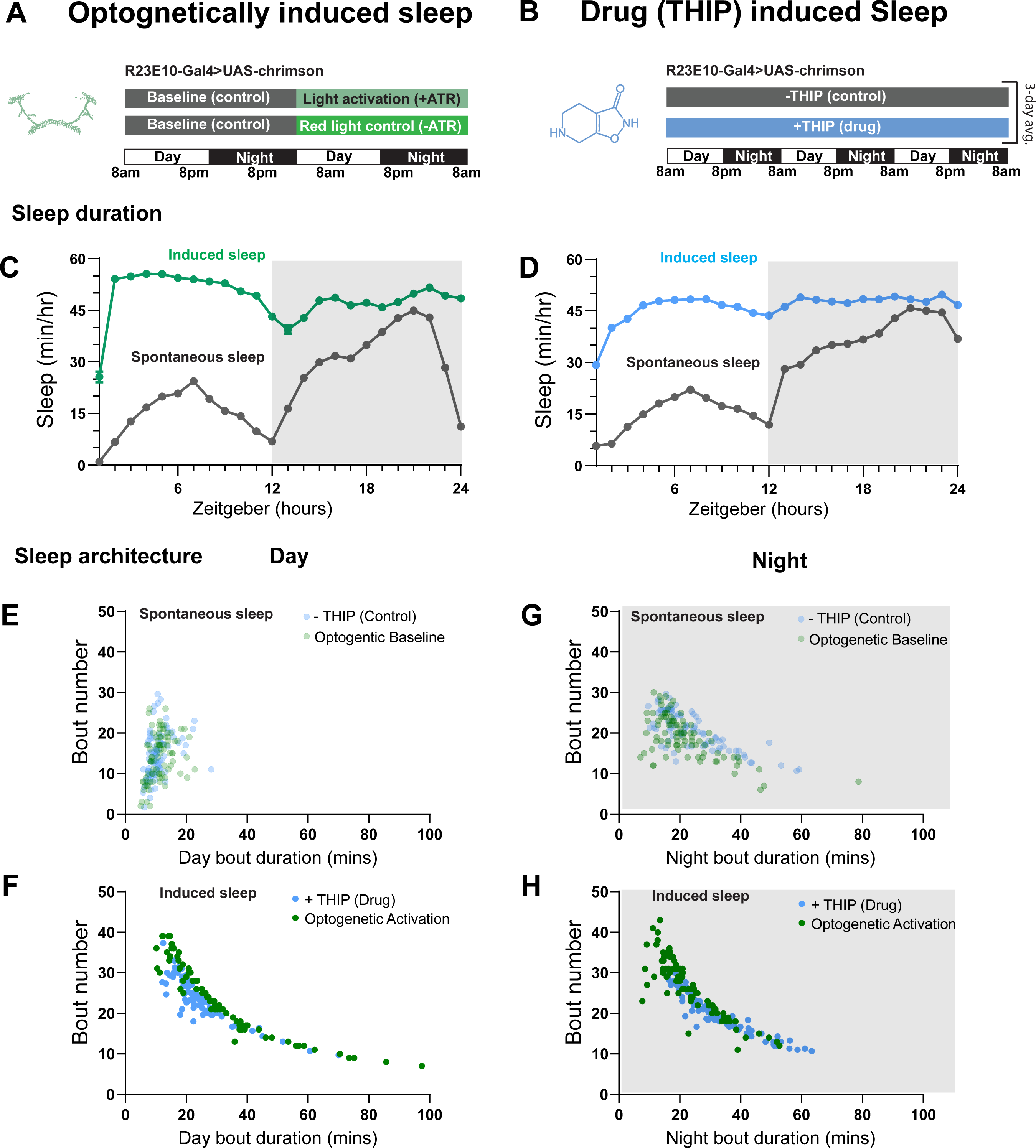
Optogenetic- and THIP-induced sleep have similar effects on sleep duration and consolidation. **A)** Experimental regime for observing the effects of optogenetic activation and THIP provision **(B)**. **C)** Sleep profile across 24 hours in the baseline condition (grey) and optogenetic activation condition (green). **D)** 3-day average of the 24-hour sleep profile of control (grey) and THIP fed (blue) flies. **E)** Daytime sleep consolidation scatterplot for optogenetic baseline and THIP control flies. **F)** Daytime sleep consolidation scatterplot for optogenetic- and THIP-induced sleep. **G)** Night-time sleep consolidation scatterplot for optogenetic baseline and THIP control flies. **H)** Night-time sleep consolidation scatterplot for optogenetic- and THIP-induced sleep. n = 87 for optogenetic activation across three replicates; n = 88 for –THIP, n = 85 for +THIP, across three replicates. Maximum bout duration possible is 720 minutes, or 12 hours. See **Supplementary File 1** for statistics.

**Figure 3.**
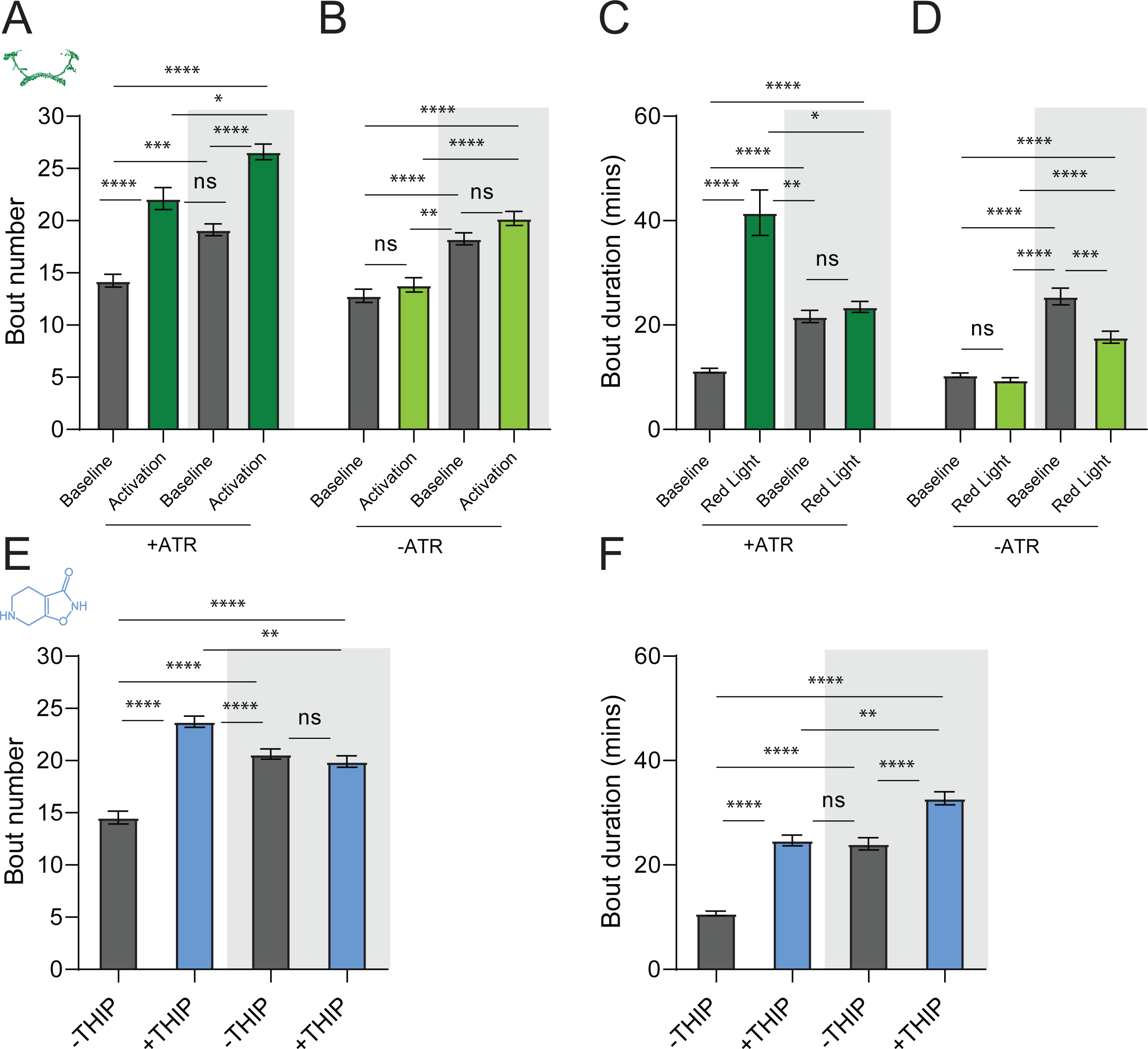
Sleep architecture in optogenetic and THIP induced sleep. **A-B.** Average number of sleep bouts in control (grey) and optogenetic activation (green) conditions in the day and night for both +ATR **(A)** and –ATR **(B)** fed flies. optogenetic-induced sleep results in an increase in the number of sleep bouts both during the day and the night, whereas red light alone has no effect. **C-D.** optogenetic activation (green) increases the average sleep duration during the day, but not the night when compared to controls (grey) in +ATR flies **(C)**. **D.** –ATR flies show no difference in mean sleep bout duration during the day, but show a decrease in average bout duration during the night. THIP (blue) increases both the average number of sleep bouts **(E)** and the average duration of sleep bouts **(F)** during the day, but not the night, when compared to controls (grey). Analysis for a and b = Kruskal-Wallis test with Dunn’s multiple comparison correction. * = p<0.05, *** = p < 0.001, **** = p<0.0001. For e and f, analysis = Ordinary one-way ANOVA with Tukey correction for multiple comparisons. *** = p<0.001, **** = p <0.0001.

## THIP-induced sleep decreases brain activity and connectivity

The brain presents an obvious place to look for any potential differences between sleep induction methods. In a previous study employing whole-brain calcium imaging in tethered flies we showed that optogenetic activation of the R23E10 circuit promotes wake-like sleep, with neither neural activity levels nor connectivity metrics changing significantly even after 15min of optogenetically-induced sleep [10]. We therefore utilized the same fly strain as in that study (R23E10-Gal4>UAS-Chrimson88-tdTomato;Nsyb-LexA>LexOp-nlsGCaMP6f) to examine the effect of THIP-induced sleep on brain activity (**Figure 4A,B**). Since we were interested in comparing acute sleep induction effects on brain activity (as opposed to prolonged sleep induction effects on behavior, as in Figures 2,3), we adapted our calcium imaging approach to allow a brief perfusion of THIP directly onto the exposed fly brain (**Figure 4A**, see Methods). As done previously for examining optogenetically-induced sleep [10], we examined calcium transients in neural soma scanning across 18 optical slices of the central fly brain (**Figure 4B**, left) and identified regions of interest (ROIs) corresponding to neuronal soma in this volume (**Figure 4B**, right, and see Methods). As shown previously [10], optogenetically activating the R23E10 circuit renders flies asleep (albeit twitchy at times) without changing the average level of neural activity measured this way (**Figure 4C**). We have previously shown that even 15min of R23E10 activation fails to produce a ‘quiet’ sleep stage that is evident after 5min in spontaneously sleeping flies [10], suggesting that this manipulation promotes ongoing active sleep until the red light is turned off – although this has not been investigated for longer epochs, e.g. the chronic activation experiments described in Figure 2. To determine the effect of THIP on neural activity in the exact same strain, we transiently perfused onto the fly brain the minimal THIP dosage required to reliably promote sleep in flies within five minutes (0.2mg/ml) [9]. In contrast to optogenetic-induced sleep, we observed overall decreased neural activity coincident with the flies falling asleep, and flies remained asleep well after the drug was washed out (**Figure 4D**). To ensure that we were actually putting flies (reversibly) to sleep in this preparation, we probed for behavioral responsiveness by puffing air onto the fly once every minute (50 ms duration, 10 psi) (**Figure 5A**). Since the time when flies fell asleep following five minutes of THIP perfusion could be variable [9], arousal probing during sleep was only initiated after 5 min of complete quiescence (**Figure 5A**, behavioral responsiveness testing). We observed decreased arousability for flies that had been induced to sleep via THIP perfusion (**Figure 5B**). Drug-induced sleep was however reversible, with flies returning to baseline levels of behavioral responsiveness to the air puffs ∼20-30 min after sleep initiation. This confirmed that the brief exposure to THIP was indeed putting flies to sleep, with an expected sleep inertia lasting the length of a typical spontaneous sleep bout [15, 16]. This contrasts with optogenetic sleep induction using the R23E10-Gal4 circuit, which does not show appear to show any sleep inertia (**Figure 4C**) [36].

**Figure 4.**
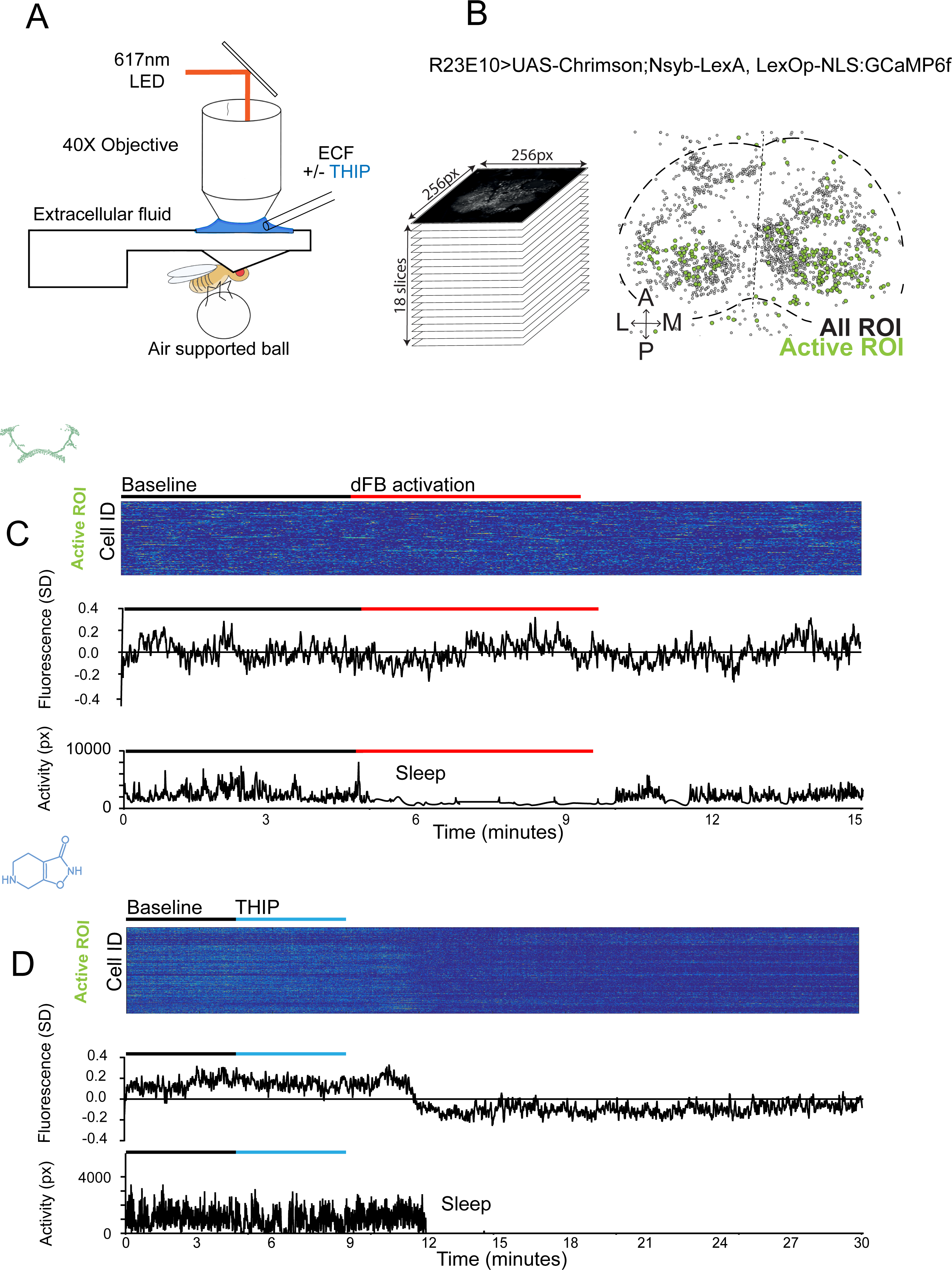
Brain imaging during optogenetic and THIP-induced sleep. **A)** Flies were mounted onto a custom-built holder that allowed a coronal visualization of the brain through the posterior side of the head. Perfusion of extracellular fluid (ECF) occurred throughout all experiments. A 617nm LED was delivered to the brain through the imaging objective during optogenetic experiments. During THIP experiments, 4% THIP in ECF was perfused onto the brain through a custom perfusion system. Behavior was recorded as the movement of flies on an air suspended ball. **B)** Left: Imaging was carried out across 18 z- slices, with a z-step of 6μm. Each z-plane spanned 667μm x 667μm, which was captured across 256 x 256 pixels. Right: A collapsed mask from one fly of neurons found to be active (green) in C alongside all identified regions of interest (ROIs, gray). **C)** Neural activity in an example fly brain, represented across cells (top) and as the population mean (middle) did not change following optogenetic-induced sleep (bottom). **D)** Neural activity in an example fly brain, represented across cells (top) and as the population mean (middle) showed an initial high level of activity in the baseline condition, which decreased when the fly fell asleep (bottom) following THIP exposure. The Y axis scale is standard deviation of the experiment mean, so the baseline is not absolute but rather reflects any difference with the overall experiment mean.

**Figure 5.**
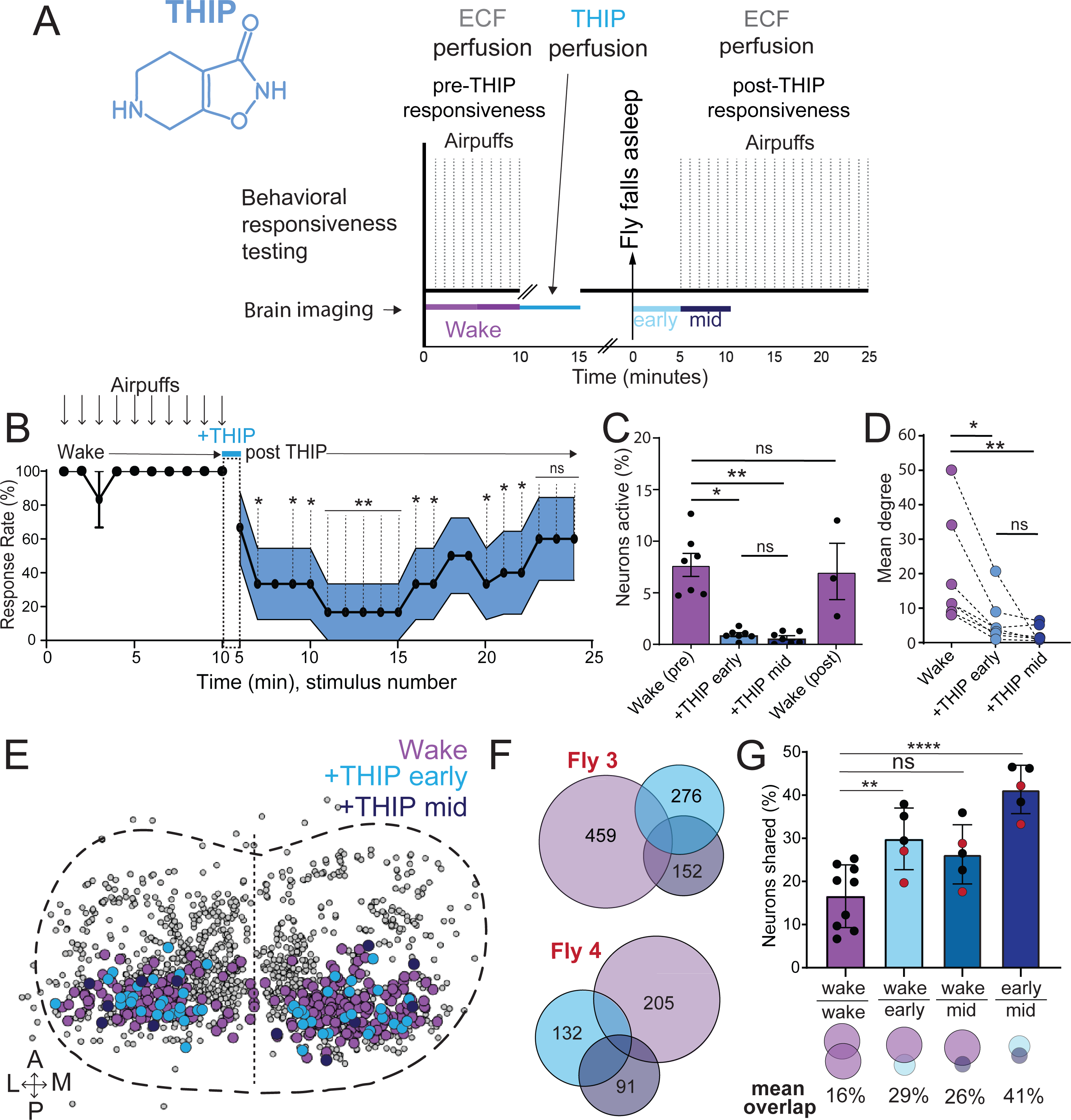
Brain activity and connectivity decreases during THIP-induced sleep. **A)** Experimental protocol for behavioral responsiveness and brain imaging experiments. 5 mins of baseline condition were recorded, during which the exposed brain was perfused with extracellular fluid (ECF), followed by 5 mins of THIP perfusion. Following sleep induction, an additional 10 minutes of calcium activity was recorded, which was separated into ‘Early’ and ‘Mid’ sleep for analysis. Air puff stimuli were delivered to test for behavioral responsiveness. **B)** Mean behavioral response rate (% ± sem) to air puff stimuli over the course of an experiment (n = 6). Air puff delivery times are indicated by the solid dots. **C)** Percent neurons active (± sem) in non ATR-fed UAS:Chrimson / X; Nsyb:LexA/+; LexOp:nlsGCaMP6f / R23E10:Gal4 flies during wake, THIP-induced sleep, and recovery (n = 9; 3 flies were recorded post-waking). **D)** Correlation analysis (mean degree ± sem) of active neurons in (C). **E)** Collapsed mask of neurons active during wake, and both early and mid THIP sleep. **F)** Overlap in neural identities between wake and THIP-induced sleep in two example flies. Number indicates active neurons within each condition; same color code as in (E). **G)** Quantification of neural overlap data. Red dots indicate the flies shown in (F). n = 9 flies. All tests are one-way ANOVA with Dunnett’s multiple comparison test. ns = not significant, * = p<0.05, ** = p<0.01, **** = p < 0.001.

We then examined more closely neural activity in flies that had been put to sleep with THIP. We found that neural activity decreased rapidly within 5 min after sleep onset (**Figure 5C**, +THIP, early). Correlation analysis also revealed a decrease in connectivity among the remaining active neurons (**Figure 5D**, +THIP, early). We also analyzed the next 5 min of sleep and observed similar results (**Figure 5C,D**, +THIP, mid). All flies eventually woke up from THIP-induced sleep, and brain activity returned to wake levels in three flies that were recorded throughout (**Figure 5C**). These observations suggest that acute THIP exposure is promoting rapid entry into a ‘quiet’ sleep stage in flies, bypassing the wake-like sleep evident during the first 5 min of spontaneous sleep onset, and closely resembling spontaneous quiet sleep typically seen after 5 minutes of inactivity [10]. Importantly, THIP-induced sleep appears to be dissimilar from optogenetically-induced sleep in this genotype, at the level of neural activity as well as connectivity [10].

In recent work we showed that rendering flies unresponsive with a general anesthetic, isoflurane, in surprising contrast to THIP, does not quieten the fly brain [36]. Isoflurane also decreased neural connectivity, as well as the diversity of neural activity across the fly brain, whereas optogenetic-induced sleep did not show any differences in neural activity or ensemble dynamics [36]. Since in our current THIP experiments we were similarly recording from neural soma that we could track through time, we were able to assess the level of overlap between the neurons that remained active during THIP-induced sleep and wakefulness (**Figure 5E, F**). We found that ∼30% of active neurons during THIP-induced sleep were also active during wake (**Figure 5F, G**). We next examined whether the same neurons remained active across successive 5min epochs during THIP-induced sleep compared to wake. We found that there is significantly more overlap between successive 5min sleep epochs (41%), compared to the waking average (**Figure 5G**), suggesting less neural turnover during THIP-induced sleep than during wake. This contrasts with optogenetic sleep, where the rate of neural turnover was not different from wake [36]. Taken together, our calcium imaging data confirm that pharmacological sleep induction promotes a different kind of sleep than optogenetic sleep induction in the same strain. Henceforth, we call this ‘quiet’ sleep, in contrast to the ‘active’ sleep that seems to be engaged by optogenetic activation [2, 10]. Notably, calcium imaging of spontaneous sleep bouts in *Drosophila* also revealed active and quiet sleep stages [10], suggesting that both of our experimental approaches are physiologically relevant. Whether drug perfusion to the brain is equivalent to feeding is of course less clear. When feeding on 0.1mg/ml THIP-laced food, flies were continuously exposed to the drug over days, with comparatively less reaching the brain. With perfusion, the brain was directly exposed to 0.2mg/ml THIP for only 5 minutes. Interestingly, in both cases this induces daytime sleep bouts which average around 25min (**Figure 3F, Figure 5B)**, the average duration of a spontaneous night-time sleep bout (**Figure 3F; Supplementary File 1**).

### Transcriptional analysis of flies induced to sleep by THIP provision

Our calcium imaging experiments suggest that different biological processes might be engaged by optogenetic-induced sleep compared to THIP-induced sleep. Additionally, we observed neural effects encompassing much of the fly brain (**Figure 5E**), as our recording approach exploited a pan-neural driver. We therefore wondered if either sleep-induction method might lead to differences in gene expression across the whole brain, and if these might highlight distinct molecular pathways engaged by either kind of sleep. To address this, we collected brains from flies that had been induced to sleep by either method, and compared the resulting transcriptomes with identically-handled control animals that had not been induced to sleep by these methods.

To control for genetic background, we again used the same R23E10-Gal4 > UAS-Chrimson flies as in our multi-day behavioral experiments and fed the flies either THIP or ATR, as in **Figure 2**. We only examined daytime sleep-induction effects for either method, as this is when we observed the greatest increase in sleep compared to controls (**Figure 2**), and previous work has shown that daytime sleep induction using either method achieves sleep functions [10, 23]. We present our THIP results first. Since THIP is a GABA-acting drug that probably affects a variety of processes in the brain aside from sleep, we also assessed the effect of THIP on flies that were prevented from sleeping (**Figure 6A**, left panel). Sleep deprivation (SD) was performed by mechanically arousing flies once every 20 seconds for the duration of the experiment, on a ‘SNAP’ apparatus [23, 37]. RNA was extracted from the brains of all groups of flies (+/- THIP, +/- SD) after 10 hours of daytime (8am-6pm) THIP (or vehicle) provision. Samples for RNA-sequencing were collected in replicates of 5 to ensure accuracy, and any significant transcriptional effects were thresholded at a log fold change of 0.58 (see Methods).

**Figure 6.**
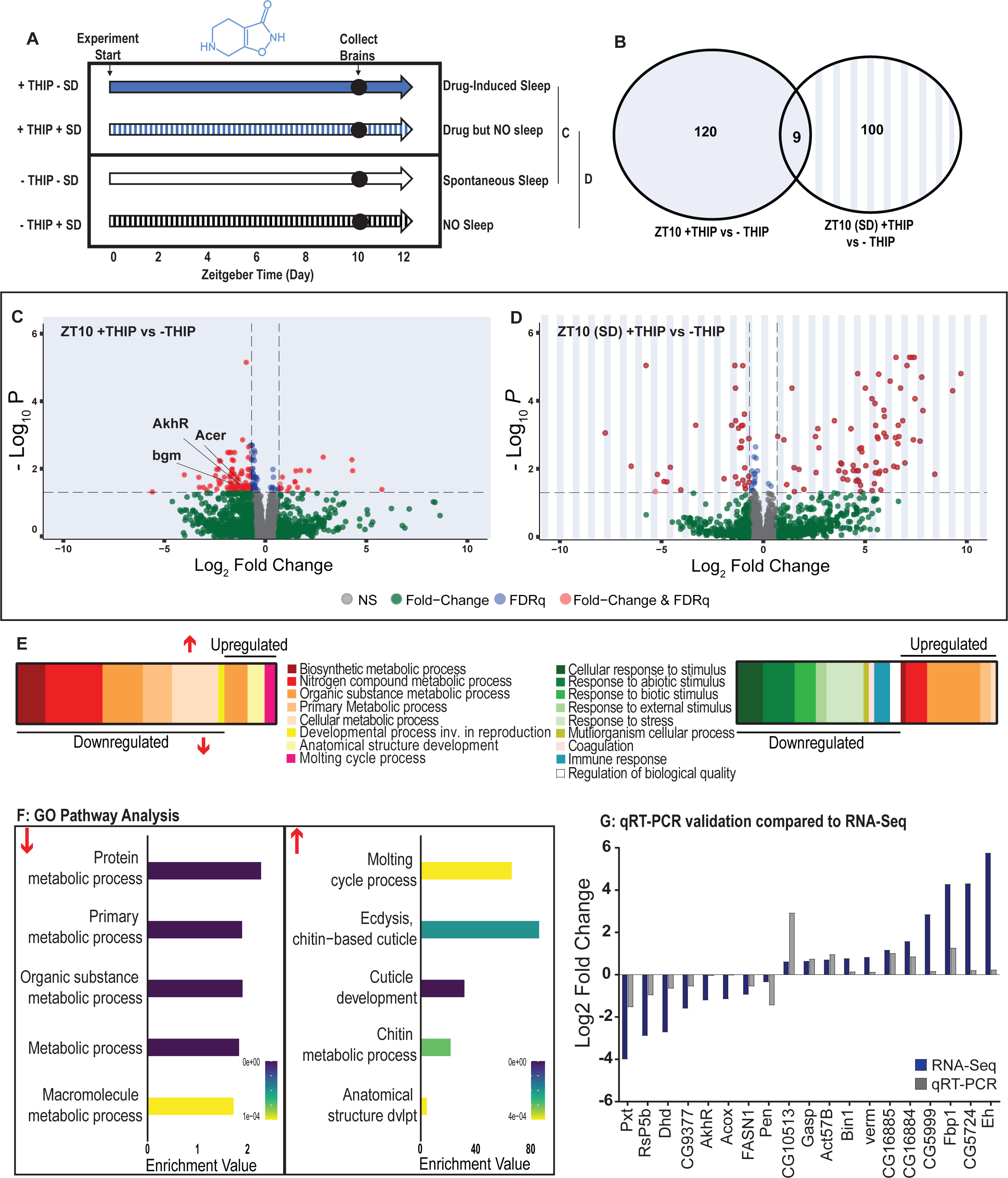
Metabolic processes are downregulated during THIP-induced sleep. **A)** Schematic representation of the experimental set-up and samples processed using RNA- Sequencing. **B)** Venn diagram showing the gene expression overlap between flies that had been treated with THIP versus their control (shaded blue) and flies that had been treated with THIP in a sleep deprived background versus their control (shaded blue bars). The number of significant differentially-expressed genes in each category is indicated. **C)** Volcano plot representing the distribution of differentially expressed genes in the presence or absence of THIP. Genes that are significantly up/down regulated meeting a Log2Fold change of 0.58 and FDRq value of 0.05 are shown in red. Genes meeting the threshold for FDRq value only are shown in blue. Fold change only is shown in green. Those genes not meeting any predetermined criteria are shown in grey. **D)** Volcano plot representing the distribution of differentially expressed genes in the presence or absence of THIP in a sleep deprived background. Criteria as above (C). **E)** Schematic representation of Gene Ontology (GO) enrichment of biological process results. Colour coded to indicate parent and child terms for comparisons between groups highlighted above (C) - Left and (D) - Right). **F)** Bar chart representation of a subset of interesting significant GO pathway terms originating from the organic substance and primary metabolic processes for the dataset shown in (C). **G)** Comparison between significant gene hits obtained via RNA-Sequencing (Blue) and qRT- PCR (Grey) in response to THIP, represented by Log2Fold change values. See **Figure 6-figure supplements 1&2, Figure 6-source data 1&2.**

Flies allowed to eat food containing 0.1mg/ml THIP *ad lib* over 10 daytime hours led to 129 significant changes in gene expression compared to vehicle-fed controls, with the large majority (110) being downregulated and only 19 upregulated (**Figure 6B,C,E; Figure 6-source data 1**). In contrast, when THIP-fed flies were prevented from sleeping this led to mostly upregulated genes (88 upregulated vs 21 downregulated, **Figure 6B,D,F; Figure 6- source data 2**), with many of the upregulated genes showing a high level of fold-change in expression. Not surprisingly, preventing sleep in THIP-fed flies led to an almost entirely non-overlapping set of gene expression changes (**Figure 6B**). This suggests that the large number of down-regulated genes in +THIP / -SD flies pertain to sleep processes rather than the effect of ingesting THIP, since only a few (9) genes overlapped with the +THIP / +SD dataset which revealed mostly upregulated genes.

Gene Ontology analysis on genes that were downregulated as a result of THIP-induced sleep highlighted a significant enrichment of metabolism pathways (**Figure 6E,F; Figure 6-figure supplement 1**). The top Gene Ontology biological processes included primary, organic substance, cellular, biosynthetic and nitrogen compound metabolic pathways, as well as ribosomal processes. Interestingly, these downregulated processes are largely consistent with a recently published mouse sleep transcriptome study [38]. Among the metabolism pathways uncovered in this dataset we observed over-representation of expected genes such as *bgm* (bubblegum CG4501) and *Acer* (Angiotensin-converting enzyme-related CG10593). Both of these genes are found in the primary metabolic and organic substance metabolic processes as well as within the Sleep Gene Ontology dataset (GO:0030431). Another downregulated metabolic gene is AkhR (adipokinetic hormone receptor), which has been found to be involved in starvation-induced sleep loss in *Drosophila* [39]. AkhR belongs to the Class A GPCR Neuropeptide and protein hormone receptors which are a gene class involved in storage fat mobilization, analogous to the glucagon receptor found in mammals [40].

Although THIP-induced sleep overwhelmingly led to gene downregulation, a few genes (19) were significantly upregulated. Gene Ontology analysis on these upregulated genes highlighted enrichment in varying groups including developmental processes and multicellular organismal processes (**Figure 6E,F; Figure 6-figure supplement 1**). Some groups were enriched under the organic substance metabolic process pathways; however, there was no overlap when comparing these to the pathways enriched due to downregulation of genes. There were some overlapping enriched pathways when we compared the gene sets from sleep-deprived flies which had also been treated with THIP (**Figure 6F; Figure 6-figure supplement 2**). However, the gene sets they involve are upregulated in the SD dataset but downregulated in sleeping flies. Interestingly, the non-sleeping THIP dataset uncovered a significant enrichment of pathways involved in the response to stress. This might be expected for flies exposed to regular mechanical stimuli over 10 hours. None of these pathways featured in the THIP sleep dataset.

To validate these findings, we conducted qRT-PCR analyses on 19 genes from our THIP sleep dataset and compared these results to our original transcriptional data. The genes represented a range of both up – and down-regulated genes, and we found convincing correspondence between the groups (**Figure 5G**). qRT-PCR comparisons with RNAseq data and associated statistics are presented in **Supplementary File 2**.

### Transcriptional analysis of flies induced to sleep by optogenetic activation

We next examined the effect of optogenetically-induced sleep on the whole-brain transcriptome, to compare to our THIP-induced sleep data. Based on our earlier findings that showed that optogenetic activation results in rapidly inducible sleep behavior that consolidates over at least 12 daytime sleep hours (**Figure 2C,E,G**), as well as our previous study showing that 10 daytime hours of optogenetic activation corrects attention defects in sleep-deprived flies [10], we induced sleep in R23E10-Gal4 x UAS-Chrimson flies for 10 daytime hours and collected tissue for whole-brain RNA-sequencing (**Figure 7A**). We selected two time points for collection, for both the sleep-induced flies (+ATR) as well as their genetically identical controls that were not fed ATR (-ATR; **Figure 7A**). Optogenetic activation was matched to the normal day-time light cycle (8 AM – 8 PM). The first collection point was after 1 hour (ZT1, 9 AM) of red-light exposure, to control for effects of ATR provision (when compared to ZT1, -ATR controls) as well as to uncover any potential short-term genetic effects of optogenetic activation. We then collected flies after 10 hours of red-light exposure (ZT 10, 6 PM) to examine longer-term genetic effects of optogenetic sleep induction, and to match exactly our THIP sleep collection timepoint (i.e., 10 hours of induced daytime sleep by either method). The combined collection points also allowed us to compare transcriptional profiles between conditions (e.g., ZT10 +ATR vs. ZT10 -ATR), to identify sleep genes, as well as within conditions (ZT1 vs. ZT10), to account for genetic effects potentially linked to light-dark cycles. As for the THIP sleep data in the same strain, samples for RNA-sequencing were collected in replicates of 5 to ensure accuracy, and any significant transcriptional effects were thresholded at a log fold change of 0.58 (see Methods).

**Figure 7.**
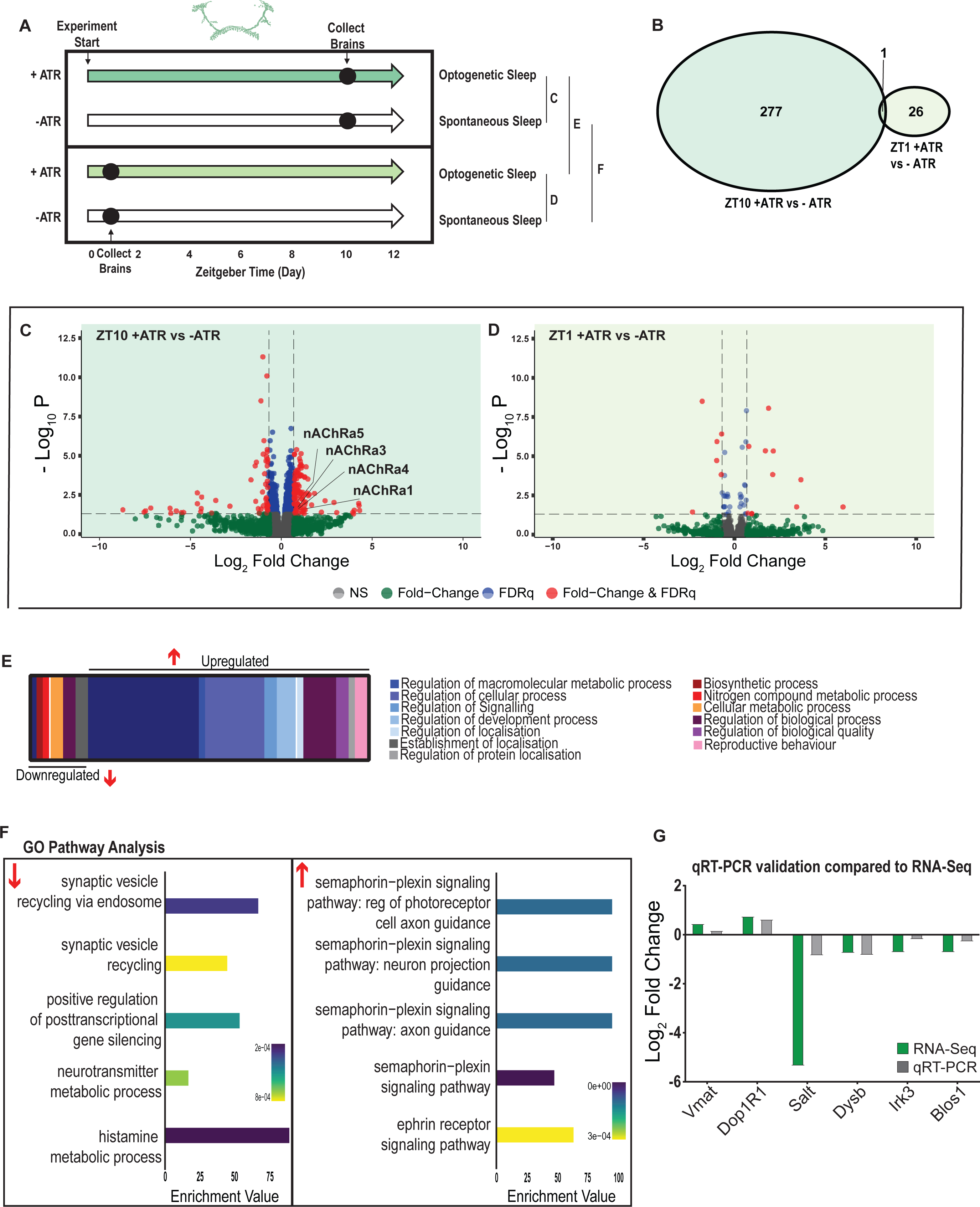
A variety of biological processes including axon guidance are upregulated during optogenetic-induced sleep. **A)** Schematic representation of the experimental set-up and samples processed using RNA-Sequencing. **B)** Venn diagram showing the gene expression overlap between flies that experienced 10 hours of optogenetic-induced sleep (ZT10) compared to -ATR controls (ZT10) and those flies where optogenetic activation was restricted to 1 hour (ZT1) and compared to -ATR controls (ZT1). **C)** Volcano plot representing the distribution of differentially expressed genes resulting from optogenetic optogenetic activation for 10 hours versus control flies which were allowed to sleep spontaneously for 10 hours. Genes that are significantly up/down regulated meeting a Log2Fold change of 0.58 and FDRq value of 0.05 are shown in red. Genes meeting the threshold for FDRq value only are shown in blue. Fold change only in green. Those genes not meeting any predetermined criteria are shown in grey. **D)** Volcano plot representing the distribution of differentially expressed genes resulting from optogenetic activation for 1 hour versus control flies which were allowed to sleep spontaneously for 1 hour. Criteria as above (C). **E)** Schematic representation of Gene Ontology (GO) enrichment of biological process results. Color coded to indicate parent and child terms comparing flies that had been activated optogenetically for 10 hours versus flies which had been allowed to spontaneously sleep for the same duration. **F)** Bar chart representation of a subset of interesting significant GO pathway terms originating from the regulation of cellular processes and signalling biological processes. **G)** Comparison between significant gene hits obtained via RNA-Sequencing (Green) and qRT-PCR (Grey) in response to optogenetic sleep, represented by Log2Fold change values. See **Figure 7-figure supplements 1-3, Figure 7-source data 1-4.**

We first examined the effect of 10 hours of daytime optogenetic-induced sleep. Here, we compared ATR-fed R23E10-Gal4 x UAS-Chrimson flies to genetically identical animals that were also exposed to red light for 10 hours but not provided with ATR in their food (ATR-). The control flies were therefore never induced to sleep by optogenetic activation, although they were still able to sleep spontaneously (see **Figure 2C,E,G**). We found that 10 hours of optogenetic activation led to 278 significant transcriptional changes, comprising mostly of upregulated genes, with 171 upregulated compared to 107 downregulated (**Figure 7B,C,E; Figure 7-source data 1**). In contrast to the THIP-induced sleep dataset, transcriptional analysis of 10hr optogenetic sleep induction uncovered a variety of different processes predominantly related to the regulation of biological and cellular processes, rather than metabolism specifically (**Figure 7E; Figure 7-figure supplement 1**). For example, of the genes that were overexpressed there is an enrichment of the Semaphorin-plexin signaling pathway (GO:0071526, GO:1902287, GO:1902285 and GO:2000305) and the ephrin receptor signaling pathway (GO:0046011), both of which are known to be involved in axonal guidance (**Figure 7F**). Interestingly, several upregulated genes code for different subunits of nicotinic acetylcholine receptors (nAchRα1,3,4 &5; **Figure 7C**). Importantly, there was almost no overlap with our sleep deprivation dataset (**Figure6-figure supplement 2; Figure6-source data 2**), suggesting that optogenetic activation is not sleep depriving (or stressing) the flies (only one upregulated gene was shared, CG40198). Of the genes that were downregulated there is enrichment of pathways that relate to synaptic vesicle recycling (GO:0036465 and GO:0036466) as well as neurotransmitter metabolic processes (GO:0042133) (**Figure 7F; Figure 7-figure supplement 1**).

In contrast to the 10hr timepoint, 1 hr of optogenetically-induced sleep had far fewer transcriptomic consequences, with only 17 genes upregulated and 10 downregulated (**Figure 7B,D**). This small number of transcriptomic changes (see **Figure 7-source data 2**) may reflect the effect of ATR feeding, rather than any genes relevant to optogenetic sleep. That 9 hours of additional optogenetic sleep increased transcriptomic changes by an order of magnitude lends confidence to the interpretation that relevant genes linked to prolonged optogenetic activation are being engaged.

To account for potential genetic effects linked to light-dark expression cycles, we compared transcriptional profiles between 10 hours of optogenetic-induced sleep to 1 hour of induced sleep. Here, we found 220 differentially regulated genes (119 upregulated and 101 downregulated) when comparing ATR-fed flies at both time points (ZT10 vs ZT1, **Figure 7- source data 3**). Since the 1-hour group was collected in the morning and the 10-hour group was collected in the evening, we expected this dataset to expose a number of circadian-regulatory genes, and this is indeed what we found (**Figure 7-figure supplement 2**). We then compared these results with a parallel ZT10 vs ZT1 experiment where flies were not fed ATR. Here we uncovered 503 differentially expressed genes (252 upregulated and 251 downregulated) when comparing flies that had not been fed ATR at both timepoints (**Figure 7-source data 4**). Importantly, there were 98 genes that overlapped between these independent Z10 vs ZT1 datasets, suggesting commonalities linked to circadian processes. Indeed, GO Pathway analysis of Biological Processes revealed a number of genes involved in the regulation of the circadian rhythm among these 98 overlapping genes, including the well-known circadian genes *period, timeless, clockwork-orange, clock and vrille*. Notably, co-factors *period* and *timeless* are both upregulated whereas *clk* is downregulated, and this is replicated in both independent datasets (**Figure 7-figure supplement 2**). This correspondence with expectations for zeitgeber or light-dark effects provides a level of confidence that our respective sleep datasets are highlighting transcriptomic changes and biological pathways relevant to either sleep induction approach. Notably, there was no overlap at all in gene expression changes between optogenetic-induced sleep and THIP-induced sleep (**Figure 6-figure supplement 1 vs Figure 7-figure supplement 1; Figure 6- source data 1 vs Figure 7-source data 1**), and the respective GO pathways analyses of biological processes are also largely non-overlapping (**Figure 7-figure supplement 3**).

To validate these findings, we compared our transcriptional results with qRT-PCR on six genes. This included the dopamine receptor Dop1R1, which regulates arousal levels [41] as well as the schizophrenia susceptibility gene dysbindin (*Dysb*), which has been shown to regulate dopaminergic function [42]. We found convincing correspondence between our qRT-PCR data and our transcriptomic data (**Figure 7G**).). qRT-PCR comparisons with RNAseq data and associated statistics are presented in **Supplementary File 2**.

### Nicotinic acetylcholine receptors regulate sleep architecture

While THIP-induced sleep caused a systemic downregulation of metabolism-related genes, the effect of optogenetic-induced sleep on gene expression was not dominated by a single category. This may be consistent with our earlier observation that brain activity looks similar to wake during optogenetic-induced sleep [10], so we could essentially be highlighting biological processes relevant to an awake fly brain, such as dopamine function [43].

However, optogenetic activation of the R23E10 circuit is not like wake, in that flies are rendered highly unresponsive to external stimuli, so perhaps like REM sleep in mammals a different category of molecular processes could be involved. In mammals, acetylcholine generally promotes wakefulness and alertness, but activity of cholinergic neurons is also high during REM sleep [44]. Neurotransmission in the insect brain is largely cholinergic, with 7 different nicotinic ‘alpha’ receptor subunits expressed in neural tissue [45]. Interestingly, four of these subunits were significantly upregulated in our optogenetic sleep dataset: nAchRα1, nAchRα3, nAchRα4, and nAchRα5 (with nAchRα6 and nAchRα7 approaching significance, **Figure 7-source data 1**). For comparison, none of these were upregulated in our sleep deprivation dataset, suggesting a sleep-relevant role. Previous studies have demonstrated a role for some of these same receptor subunits in sleep regulation, in particular nAchRα4 (also called *redeye*) which is upregulated in short-sleeping mutants [46] and nAchRα3 which has been reported to regulates arousal levels in flies [47]. Together, these studies suggest processes that might be reconsidered in the context of active sleep, as highlighted by our gene expression findings. We therefore sought to examine the role of cholinergic signalling in sleep more closely, by knocking out each nAchRα subunit and examining effects of each subunit knockout on sleep architecture. Since our transcriptomic analysis encompassed effects of active sleep on whole-brain gene expression, we first eliminated each nAchRα subunit across the brain, by testing confirmed genetic deletions [48, 49].

We first examined the effect of each nAchRα subunit deletion on sleep duration, using the 5- minute criterion for quantifying sleep in *Drosophila* [15]. We found that the nAchRα mutants fell into two different categories: removal of nAchRα1 and nAchRα2 significantly decreased sleep, day and night; whereas removal of nAchRα3, nAchRα4, nAchRα6 and nAchRα7 significantly increased sleep, day and night (**Figure 8A**). The nAchRα5 knockout was found to be homozygous lethal, so was not included. To examine sleep architecture in these mutants, we quantified sleep bout number and duration and plotted these together as done previously for our sleep induction experiments (**Figure 2**). Examining the data this way, it is clear to see how nAchRα1 and nAchRα2 are different: most sleep bouts are very short, day and night (**Figure 8B**, top 2 rows, left panels, green dots). In contrast, knocking out the other alpha subunits seems to consolidate sleep, especially at night (**Figure 8B**, bottom 4 rows, left panels). nAchRα3 knockouts were most striking in this regard, with these flies sleeping uninterrupted for an average of 156.53 minutes (± 18.06) during the day and 160.76 minutes (± 17.92) at night. Increased sleep consolidation in these mutants was however not due to lack of activity. While awake, nAchRα3 animals were just as active as controls (activity per waking minute = 2.69 ± 0.18 versus 2.5 ± 0.06, respectively). However, some knockouts did increase waking activity levels; waking activity data for all of the knockout strains and their genetic controls are presented in **Figure 8-figure supplement 1**.

**Figure 8.**
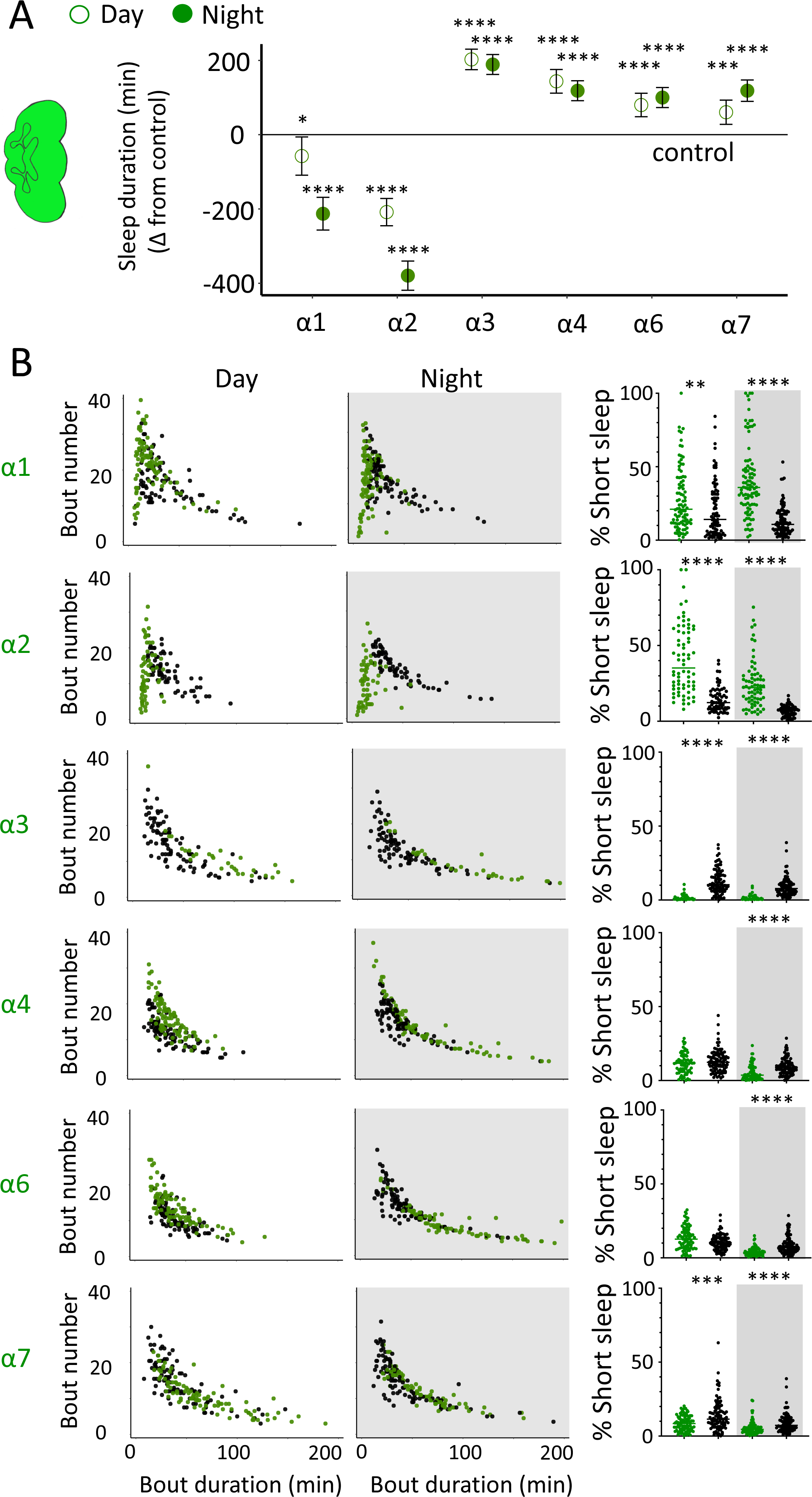
nAchRα subunit knockouts bidirectionally regulate >5min sleep as well as short sleep. **A.** Average total day and night sleep duration (minutes±95% confidence intervals) in nAchRα knockout mutants, expressed as difference to their respective background controls (see Methods). α1, N=91; contro1 (X^59^w^1118^) = 93; α2, N=70; control (w^1118^ActinCas9) = 65; α3, N=43; (ActinCas9) =9; α4, N=87; (w^1118^ActinCas9) =98; α6, N=91; (w^1118^ActinCas9) =91; α7, N=94; (ActinCas9) =95. \**P*<0.05, \*\*\**P*<0.001, \*\*\*\**P*<0.0001 by t-test adjusted for multiple comparisons. **B.** Left two panels: sleep architecture for the same six knockout strains as in A (green), shown against their respective controls (black). Each datapoint is a fly. Right panels: cumulative short sleep (1-5min) expressed as a percentage of total sleep duration. Data are the from the same experiment as in A&B. Each datapoint is a fly. \*\**P*<0.01, \*\*\**P*<0.001, \*\*\*\**P*<0.0001 Man-Whitney U Test. All data were collected over three days and three nights and averaged.

We next questioned what kind of sleep the nAchRα knockout flies might be getting. In previous work we have shown that flies can be asleep already after the first minute of spontaneous inactivity, and that during the first five minutes of sleep the fly brain displays wake-like levels of neural activity [10]. We have termed this early sleep stage ‘active sleep’ to distinguish it from ‘quiet sleep’ that typically follows after 5-10 minutes [50]. One way of estimating the amount of ‘active sleep’ in *Drosophila* flies is to sum all short sleep epochs lasting between 1-5 minutes and expressing this as a percentage of total sleep [10]. When we re-examined our nAchRα knockouts in this way, we found that this behavioral readout for ‘active sleep’ was significantly affected by the loss of select nAchRα subunits. Short sleep increased significantly during both the day and the night in nAchRα1 and nAchRα2 (**Figure 8B**, top 2 rows, right panel, green dots). In contrast, and consistent with our sleep architecture analyses (above), nAchRα3 displayed almost no short sleep (**Figure 8B**, row 3, right panel). Finally, in nAchRα4 and nAchRα6 short sleep was significantly decreased at night, while in nAchRα7 short sleep was significantly decreased day and night (**Figure 8B**, rows 4-6, right panel). In conclusion, every one of the nAchRα knockouts we tested affect short sleep in some way, either increasing (nAchRα1 and nAchRα2) or decreasing it (nAchRα3, nAchRα4, nAchRα6, nAchRα7). Additionally, tallying the proportion of short sleep bouts provides valuable insight into altered sleep architecture in mutant strains.

We nevertheless questioned whether these systemic effects of nicotinic receptors on short sleep were perhaps a trivial consequence of altered >5min sleep duration in these mutants, especially regarding the striking differences between nAchRα1&2 and the other subunit knockouts. We therefore returned to our ‘quiet’ sleep (THIP) dataset to contrast a gene derived from that study. We had found that several of the THIP-induced sleep genes are involved in metabolic processes, which are mostly downregulated (**Figure 6-figure supplement 1; Figure 6-source data 1**). This included the adipokinetic hormone receptor (AkhR), which has previously been associated with starvation-induced sleep regulation [39].

We employed an RNAi strategy to downregulate this metabolic gene’s expression across the fly brain in AkhR-RNAi / R57C10-Gal4 flies (see Methods). We found that downregulating AkhR significantly decreased sleep duration during the day as well as night, compared to genetic control strains (**Figure 9A,B**). Accordingly, sleep bout duration and number decreased, especially during the day (**Figure 9C**). However, in contrast to knocking out nAchRα1 and nAchRα2, which also significantly decreased sleep duration day and night, short sleep was not significantly altered in AkhR knockdown animals compared to genetic controls (**Figure 9D**). This suggests that short (1-5min) sleep might be under separate regulatory control than >5min sleep.

**Figure 9.**
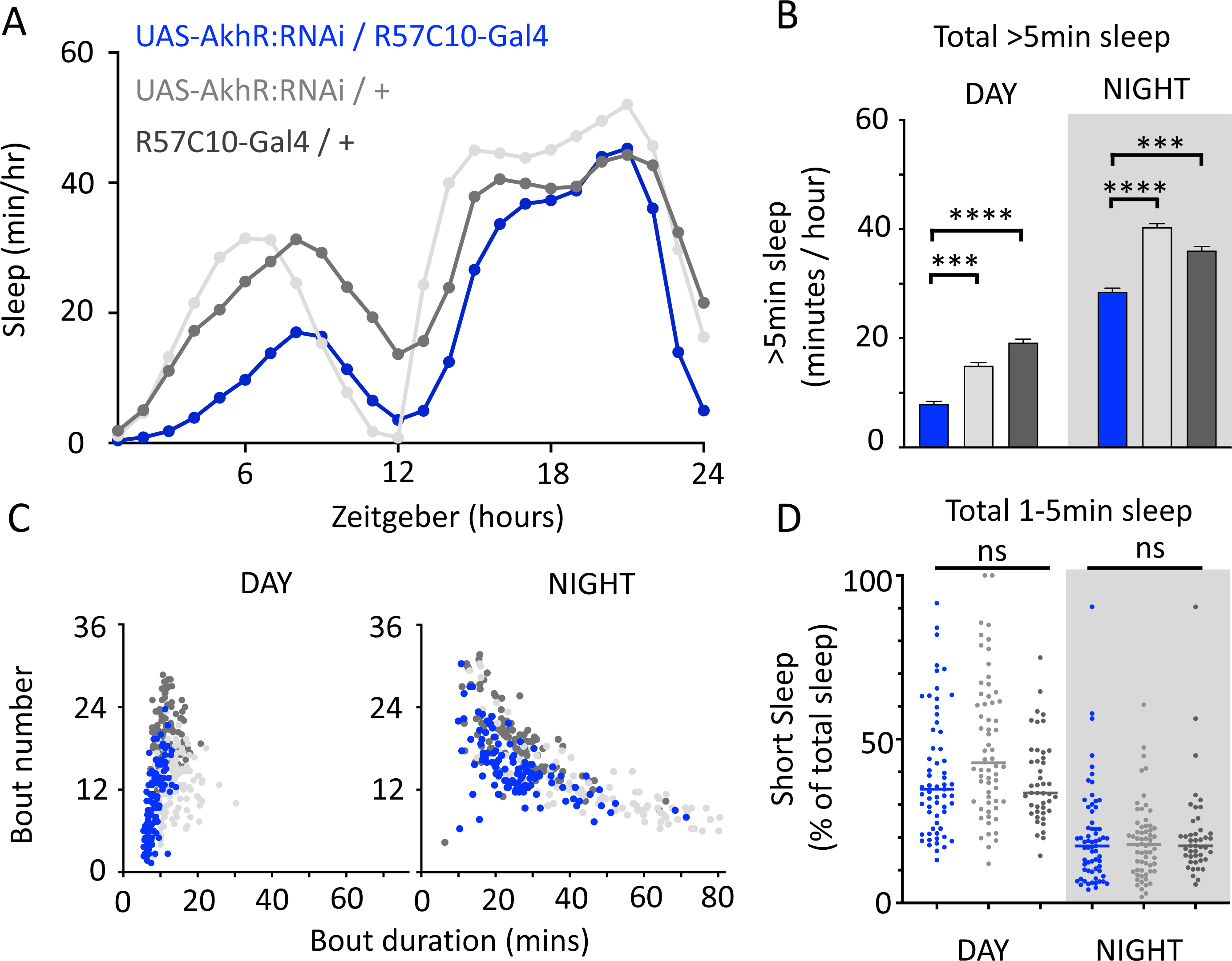
AkhR knockdown decreases >5min sleep but not short sleep. **A,B.** Total sleep (>5min) in UAS-AkhR:RNAi / R57C10-Gal4 flies (blue, N=126) compared to genetic controls (light grey: UAS-AkhR:RNAi / +, N= 124; dark grey: R57C10-Gal4 / +, N= 120). **C.** Sleep architecture (average bout duration versus bout number per fly) in data from A,B. **D.** Cumulative short sleep (1-5min, expressed as a % of total sleep) in UAS-AkhR:RNAi / R57C10-Gal4 flies (blue) compared to genetic controls (light grey: UAS-AkhR:RNAi / +; dark grey: R57C10-Gal4 / +). Wild-type background (+) is Canton-S(w^1118^). Each datapoint is a fly. \*\*\**P*<0.001, \*\*\*\**P*<0.0001 Man-Whitney U Test. ns, not significant. All data were collected over two days and two nights and averaged.

We then returned to the nAchRα subunits and employed an RNAi strategy to knock them down in the R23E10 sleep-promoting neurons specifically. While it is unlikely that these cholinergic receptor subunits are restricted to just these neurons, we were curious if removing any of them from this specific circuit had similar effect on sleep as in the knockout strains. We found that nAchRα1 knockdown in the R23E10 neurons significantly decreased total sleep during both day and night (**Figure 10A,B**), which was also true for the nAchRα1 knockouts (**Figure 8A**). Also consistent with the knockouts, the proportion of short sleep bouts was increased in this this specific knockdown (**Figure 10C**). Among all of the other alpha subunits, knockdown of nAchRα5 and nAchRα6 in the R23E10 neurons significantly increased daytime and night-time sleep, respectively (**Figure 10A,B**), while short sleep was correspondingly decreased only in nAchRα5 (**Figure 10C**). None of the other alpha subunits had any effect in these sleep-promoting neurons. This suggests that their sleep-regulatory role lies outside of this circuit, while nAchRα1, nAchRα5, and nAchRα6 might regulate sleep-related functions within this circuit.

**Figure 10.**
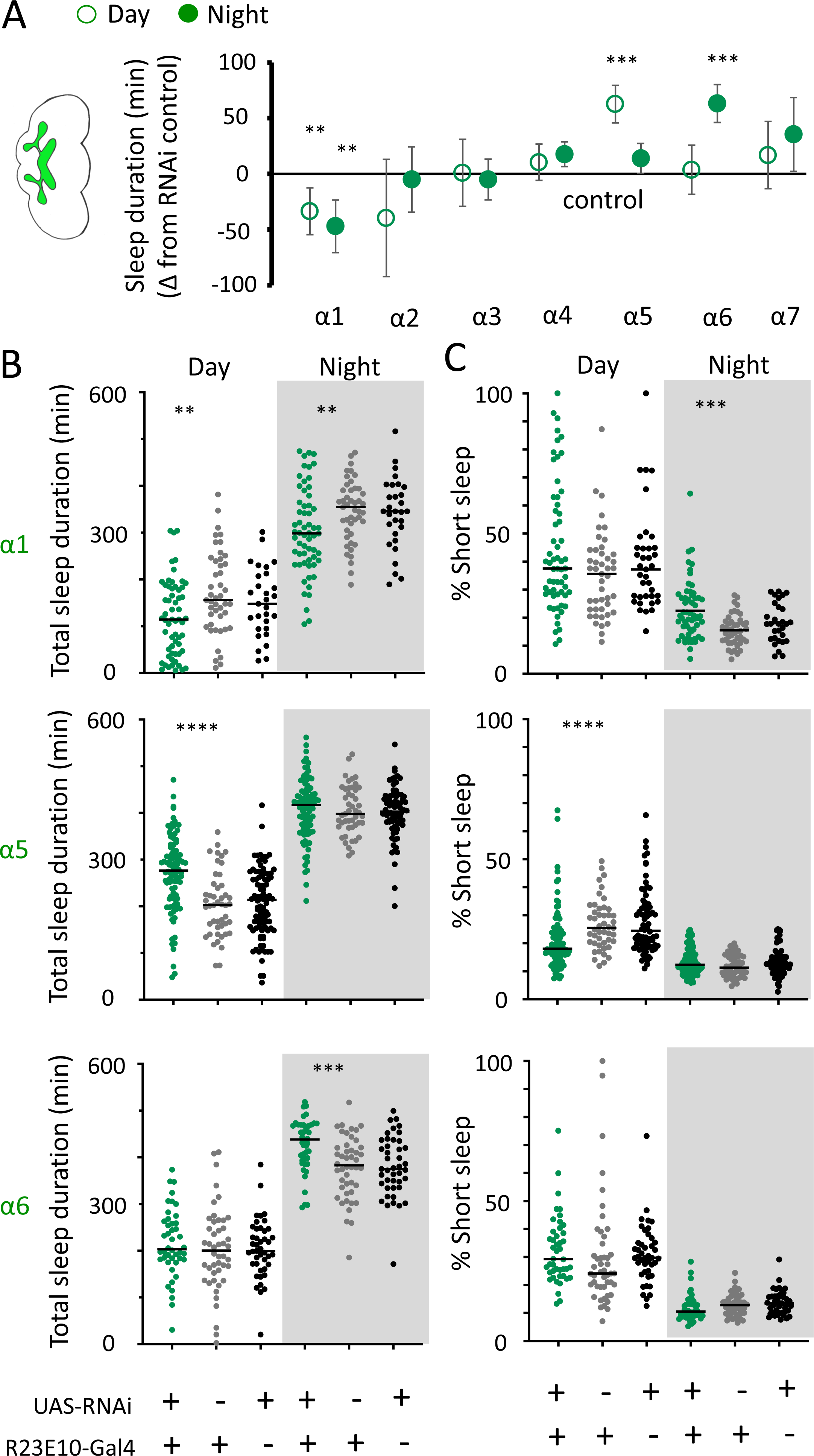
nAchRα subunit knockdowns in sleep-promoting neurons. **A.** Average total day and night sleep duration (minutes±95% confidence intervals) in UAS-nAchRα RNAi / R23E10-Gal4 flies, expressed as difference to RNAi / + controls (all were also compared Gal4 / + controls, not shown). α1 N=60, Gal4 N=45, RNAi N=37; α2 N=21, Gal4 N=24, RNAi N=50; α3 N=33, Gal4 N=50, RNAi N=59; α4 N=92, Gal4 N=90, RNAi N=80; α5 N=94, Gal4 N=47, RNAi N=74; α6 N=44, Gal4 N=47, RNAi N=43; α7 N=32, Gal4 N=57, RNAi N=44. \*\**P*<0.01, \*\*\**P*<0.001 by t-test adjusted for multiple comparisons. **B.** Total >5min sleep duration data for significant knoockdowns in (A). **C.** Cumulative short sleep (1- 5min) expressed as a percentage of total sleep duration. Data are the from the same experiment as in A&B. Each datapoint is a fly. \*\*\**P*<0.001, \*\*\*\**P*<0.0001 Man-Whitney U Test. All data were collected over two days and two nights and averaged.

## Discussion

One of the key advantages of studying sleep in *Drosophila* is that this versatile model provides a variety of reliable approaches for inducing and controlling sleep. By being able to induce sleep on demand, either genetically or pharmacologically, researchers have been able to manipulate sleep as an experimental variable, and in this way be better able to assess causality when probing potential sleep functions. However, this approach has often sidestepped the question of whether different sleep induction methods are equivalent, or whether distinct forms of sleep might be engaged by different genetic or pharmacological treatments. In mammals, GABA agonists typically promote slow-wave sleep (SWS), which has been associated with cellular homeostasis and repair process in the brain [4, 51]. In contrast, drugs targeting acetylcholine receptors, such as carbachol, have been found to promote brain states more reminiscent of REM sleep [52, 53]. Although these drugs all induce sedative (or dissociative) states, they are clearly producing dissimilar forms of sleep in mammals, with likely different functions or consequences for the brain. In *Drosophila*, evidence suggests that the GABA agonist THIP promotes a form of deep or ‘quiet’ sleep, which may be functionally analogous to mammalian SWS [9, 27, 54]. In contrast, optogenetic activation of the dFB as well as other neurons may promote a form of active sleep, which we have suggested could be achieving some similar sleep functions as REM [2]. THIP-induced sleep in flies has been associated with waste clearance from the brain [27], just as SWS has been associated with clearance of waste metabolites via the mammalian brain’s glymphatic system [51]. Such functional homology suggests that the transcriptional changes we uncovered for THIP-induced sleep in *Drosophila* might also be relevant for mammalian SWS, with these largely centered on reduced metabolic processes and stress regulation [38]. In contrast, except for the upregulation of cholinergic signaling [55], there is little to compare to test hypotheses potentially linking active sleep in flies with REM sleep in mammals, except for the potential upregulation of cholinergic signaling. Even this cholinergic connection seems odd, seeing that the predominant excitatory neurotransmitters are reversed in the brains of insects and mammals: glutamate in mammals and acetylcholine in insects [56]. Additionally, only nicotinic receptor subunits were identified in our analyses, whereas muscarinic receptors have been more commonly associated with REM sleep in mammals [57, 58]. Nevertheless, it is clear from our results that optogenetic-induced active sleep upregulates the expression of multiple nicotinic acetylcholine subunits, and that knocking these out individually has profound (and opposing) effects on sleep architecture in flies, and that a subset might function intrinsically within sleep promoting neurons. This supports other studies showing a role for nAchRα’s in sleep regulation [46, 47]. It will be especially interesting in future brain imaging studies to see whether a knockout such as nAchRα3 is eliminating one kind of sleep (e.g., active sleep) as predicted by our behavioral data, and whether this is associated with any functional consequences. Similarly, it will be telling to see whether the opposite sleep phenotypes observed in nAchRα1 for example result in a distinct class of functional consequences. A previous study has shown that nAchRα1 knockout animals have significantly shorter survivorship compared to controls, with flies dying almost 20 days earlier [59]. One reason could be because of impaired or insufficient deep sleep functions (e.g., brain waste clearance [27]). The nAchRα knockouts provide an opportunity to further examine mutant animals potentially lacking either kind of sleep, although this will have to be confirmed by brain imaging or electrophysiology.

We found little similarity between two different approaches to inducing sleep in flies, at the level of gene expression as well as acute effects on brain activity. It may however not be surprising that these entirely different sleep induction methods produce dissimilar physiological effects. After all, one method requires flies to ingest a drug which then must make its way to the brain, while the other method directly activates a subset of neurons in the central brain, as well as in the ventral nerve cord [60]. Yet both methods yield similarly increased sleep duration profiles and consolidated sleep architecture (**Figure 2**). One underlying assumption with focusing on sleep duration as the most relevant metric for understanding sleep function in *Drosophila* is that sleep is a unitary phenomenon in the fly model, meaning that primarily one set of functions and one form of brain activity are occurring when flies sleep. There is now substantial evidence that this is unlikely to be true, and that like other animals flies probably also experience distinct sleep stages that accomplish different functions [9, 10, 27, 50, 61, 62]. This does not mean that these functions are mutually exclusive; for example, both THIP provision and neuronal activation of central brain circuits have been found to promote memory consolidation in *Drosophila* [23]. Indeed, it seems reasonable to propose that different sleep stages could be synergistic, accomplishing a variety of homeostatic functions that might be required for adaptive behaviors in an animal. Our results suggest that THIP provision promotes a ‘quiet sleep’ stage in flies, which induces a brain-wide downregulation of metabolism-related genes. This is consistent with studies in flies showing that metabolic rate is decreased in longer sleep bouts, especially at night, and that this is recapitulated by THIP-induced sleep [63]. Our findings also align with a previous study showing that mitochondrial metabolic processes are upregulated in the absence of sleep [64]. One argument for why metabolism-related genes are downregulated during THIP-induced sleep might be that flies are starved (because they are sleeping more, and thus perhaps not eating). However, flies induced to sleep by optogenetic activation are also sleeping more (indeed, their average sleep bouts are longer), yet these did not reveal a similar downregulation of metabolic processes. Another view might be that our sampling was done after flies had achieved 10 hours of induced sleep, so sleep functions might have already been achieved by that time. Thus, we might not be uncovering genes required for achieving ‘quiet’ sleep functions as much as identifying exactly the opposite: genetic pathways that have been satisfied by 10 hours of induced quiet sleep. Other studies using THIP to induce sleep have examined longer timeframes (e.g., 2 days [23]), so it remains unclear whether changes in gene regulation relate to sleep functions that have been achieved or that are still being engaged. For our transcriptomic study, both sleep induction protocols were conducted during the day (thus in presumably well-rested flies) to deliberately avoid any sleep pressure confounds. Since the dFB has been proposed as a regulator of sleep homeostasis [20-22], one could question why its prolonged activation did not incur a deep sleep debt – which might have been detectable at a transcriptomic level by some overlap with our sleep deprivation controls. That we did not see this suggests induced daytime active sleep might not accrue any sleep debt. We have shown previously that daytime sleep is qualitatively different than night-time sleep in flies, which is ‘deeper’ [50]. It is for example possible that most quiet sleep happens at night in flies, so a daytime regime of induced active or quiet sleep might not incur any sleep debt for either. Following this logic, our RNA analyses simply revealed the transcriptomic changes resulting from either manipulation in well-rested flies, with the key result being that there are zero shared genes after 10 daytime hours of either manipulation.

In contrast to THIP, our optogenetic manipulation induced a brain-wide upregulation of a variety of neural mechanisms unrelated to metabolic processes, suggesting a different kind of sleep was engaged. Although many studies have shown that the R23E10-Gal4 circuit is sleep promoting (e.g.[19, 21, 62]), and we have shown that acute activation of these neurons produces a form of active sleep [10], it seems unlikely that active sleep regulation is limited to the dFB neurons alone [60]. Other circuits in the fly central brain are also sleep-promoting, including in the ellipsoid body [65] and the ventral fan-shaped body (vFB) [66], although it remains unknown if activation of these other circuits also promotes an active sleep stage, or whether a similar transcriptome might be engaged by these alternate approaches to optogenetic sleep induction in flies. This again highlights a variant of the same problem we have uncovered in the current study comparing pharmacology with optogenetics: different circuit-based approaches could all be increasing sleep duration but achieving entirely different functions by engaging distinct transcriptomes and thus different sleep functions. How many different kinds of functions are engaged by sleep remains unclear: is it roughly two functional categories linked to quiet and active sleep, or is it a broader range of sub-categories that are not so tightly linked to these obviously different brain activity states?

A compelling argument could nevertheless be made for two kinds of sleep in most animals, with two distinct sets of functions [1, 54]. Most animals have been shown to require a form of ‘quiet’ sleep to ensure survival, suggesting that these might encompass an evolutionarily conserved set of cellular processes that promote neural health and development [67], and that operate best during periods of behavioral quiescence. Nematode worms thus experience a form of quiet sleep when they pause to molt (‘lethargus’) into a different life stages during their development [68], or when cellular repair processes are needed following environmental stress [69]. In flies, quiet sleep seems to be similarly required for neuronal repair [26] or waste clearance [27], and there is evidence that glia might play a key role in these cellular homeostatic processes in flies [26] as well as other animals [70]. Thus, SWS in mammals and birds might present a narrow neocortical view of a more ancient set of sleep functions centered on quiescence and decreased metabolic rate. Indeed, neural quiescence is also a feature of SWS, both at the level of pulsed inhibition (down-states), as well as in other parts of the brain beyond the cortex [1]. Similar to findings in flies that are induced into a quiet sleep stage with THIP [63], metabolic rate also decreases during SWS in mammals [71]. In contrast, metabolic rate is similar to waking during REM sleep in mammals [72], suggesting an alternate set of functions not linked to cellular homeostasis. What might these functions be, and could some of these be conserved between active sleep in invertebrates and REM sleep in mammals? A REM-like sleep stage has now been identified in a variety of invertebrate species, including cephalopods [12] and jumping spiders [73]. In humans, REM sleep has been implicated in emotion regulation [74], and cognitive disorders where emotions are dysregulated, such as depression, are often associated with REM sleep dysfunction [75]. While it is not evident how to study emotions in insects (but see [76]), it could be argued that arousal systems more generally are employed to detect prediction errors and thereby promote learning [77]. Thus, we and others have suggested that REM sleep might be crucial for optimizing prediction, and attention, and learning [2, 54, 78], and this may involve different kinds of homeostatic mechanisms centered on brain circuits rather than cells. Whether active sleep in flies plays a similar homeostatic role remains to be determined. Our finding that optogenetically-induced sleep in *Drosophila* upregulates different nAchRα subunits is consistent with new findings that these subunits regulate appetitive memories in flies [79] and that cholinergic systems more generally underpin learning and memory in this animal [80]. Yet learning and memory in flies clearly also benefits from quiet sleep, as evidenced by multiple studies using THIP as a sleep-promoting agent [23, 25, 81]. One view consistent with our findings and previous studies is that both kinds of sleep are crucial for optimal behavior: quiet sleep for cellular homeostasis and active sleep for circuit-level homeostasis. Manipulating these separately, alongside the non-overlapping pathways engaged by either kind of sleep, should help further disambiguate the functions potentially associated with these distinct sleep stages.

## Materials and Methods

### Animals

*Drosophila melanogaster* flies were reared in vials (groups of 20 flies / vial) on standard yeast-sugar-agar based media (1.0-1.5-0.5 gram ratio) under a 12:12 light/dark (8 AM:8 PM) cycle and maintained at 25°C with 50% humidity. Adult, 3-5 day-old female, flies were used for all experiments and randomly assigned to experimental groups. Fly lines used for behavioral and RNA-sequencing experiments include R23E10-Gal4 (attp2; Bloomington 49032; Bloomington Drosophila Stock Center, Bloomington, Indiana) and UAS-CsChrimson-mVenus (attp18; Bloomington 55134; Provided by Janelia Research Campus, Ashburn, Virginia)[31]. For all 2-photon experiments, flies with the genotype 10XUAS- Chrimson88-tdTomato (attp18) / +: LexAop-nlsGCaMP6f (VIE-260b; kindly provided by Barry J. Dickson) / +: Nsyb-LexA (attP2) [82], LexAop-PAGFP (VK00005) / R23E10-Gal4 were used. Optogenetically-manipulated fly lines were maintained on food containing 0.5mg/ml all-trans retinal (ATR; Merck, Darmstadt, Germany) for 24 hours prior to assay to allow for sufficient consumption. Pharmacologically-manipulated flies were maintained on food with 0.1 mg/ml of Gaboxadol (4,5,6,7-tetrahyrdoisoxazolopyridin-3-ol, THIP) for the duration of behavioral experiment [23].

### 2-photon imaging

2-photon imaging was performed as described previously [10] using a ThorLabs Bergamo series 2 multiphoton microscope. Fluorescence was detected with a High Sensitivity GaAsP photomultiplier tube (ThorLabs, PMT2000). GCaMP fluorescence was filtered through the microscope with a 594 dichroic beam splitter and a 525/25nm band pass filter.

For imaging experiments, flies were secured to a custom-built holder [10]. Extracellular fluid (ECF) containing 103 NaCl, 10.5 trehalose, 10 glucose, 26 NaHCO3, 5 C6H15NO6S, 5 MgCl2 (hexa-hydrate), 2 Sucrose, 3 KCl, 1.5 CaCl (dihydrate), and 1 NaH2PO4 (in mM) at room temperature was used to fill a chamber over the head of the fly. The brain was accessed by removing the cuticle of the fly with forceps, and the perineural sheath was removed with a microlance. Flies were allowed to recover from this for one hour before commencement of experiments. Imaging was performed across 18 z-slices, separated by 6µm, with two additional flyback frames. The entire nlsGCaMP6f signal was located within a 256 x 256px area, corresponding to 667 x 667µm. Fly behavior was recorded with a Firefly MV 0.3MP camera (FMVU-03MTM-CS, FLIR Systems), which was mounted to a 75mm optical lens and an infrared filter. Camera illumination was provided by a custom-built infrared array consisting of 24 3mm infrared diodes. Behavioral data was collected for the duration of all experiments.

For THIP experiments, an initial five minutes of baseline activity was captured, followed by perfusion of 0.2mg/ml THIP in ECF onto the brain at a rate of 1.25ml/minute for five minutes. An additional twenty minutes of both brain and behavioral activity were recorded to allow visualization of the fly falling asleep on the ball as a result of THIP exposure. All flies were removed from imaging after an experiment and confirmed to have awoken by visual inpection. A subset of THIP-exposed flies (n=3) remained in the imaging setup to measure brain activity upon recovery.

### Behavioral responsiveness probing

For probing behavioral responsiveness in the brain imaging preparation, flies walking on an air-supported ball were subjected to a 50ms long, 10psi air puff stimulus, which was generated using a custom-built apparatus and delivered through a 3mm-diameter tube onto the front of the fly. Flies were subjected to 10 pre-THIP stimuli at a rate of one puff/minute, to characterize the baseline response rate. Flies were then perfused with 0.2mg/ml THIP in ECF for five minutes, followed by continuous ECF perfusion for the remaining experimental time. Five minutes after the fly had fallen asleep on the ball, a further 20 air puff stimuli were delivered, at a rate of one puff/minute. Behavioral responses to the air puff were noted as a ‘yes’ (1) or ‘no’ (0), which were characterized as the fly rapidly walking on the ball immediately following the air puff. For statistical analysis, the pre-THIP condition was compared to either the first or last 10 minutes of the post-THIP condition.

### Imaging analysis

Preprocessing of images was carried out using custom written Matlab scripts and ImageJ. Motion artifacts of the images were corrected as described previously [10]. Image registration was achieved using efficient sub-pixel image registration by cross-correlation. Each z-slice in a volume (18 z-slices and 2 flyback slices) is acquired at a slightly different time point compared to the rest of the slices. Hence to perform volume (x,y,z) analysis of images, all the slices within a volume need to be adjusted for timing differences. This was achieved by using the 9^th^ z-slice as the reference slice and temporal interpolation was performed for all the other z-slices using ‘sinc’ interpolation. The timing correction approach implemented here is conceptually similar to the methods using in fMRI for slice timing correction.

For each individual z-slice, a standard deviation projection of the entire time series was used for watershed segmentation with the ‘Morphological segmentation’ ImageJ plugin [83]. Using a custom-written MatLab (Mathworks) code, the mean fluorescent value of all pixels within a given ROI were extracted for the entire time series, resulting in a n x t array for each slice of each experiment, where n refers to the number of neurons in each Z-slice, and t refers to the length of the experiment in time frames. These greyscale values were z-scored for each neuron, and the z-scored data was transformed into a binary matrix where a value of > 3 standard deviations of the mean was allocated a ‘1’, and every value < 3 standard deviations was allocated a ‘0’. To determine whether a neuron fired during the entire time series, a rolling sum of the binary matrix was performed, where ten consecutive time frames were summed together. If the value of any of these summing events was greater or equal to seven (indicating a fluorescent change of > 3 standard deviations in 7/10 time frames), a neuron was deemed to be active. For THIP sleep experiments, the five minutes of inactivity occurring after an initial 30 seconds of behavioral inactivity were used. After identifying firing neurons for each condition (wake vs sleep), the percentage of active neurons was calculated in each slice by taking the number of active neurons and dividing it by the total number of neurons.

Traces of active neurons were used to calculate the number of firing events. This was done using the ‘findpeaks’ matlab function on the zscored fluorescent traces, with the parameters ‘minpeakheight’ of 3, and ‘minpeakdistance’ of 30. Data resulting from this was crosschecked by taking the binary matrices of the time traces and finding the number of times each neuron met the activity threshold described above. Graph-theory analyses of neural connectivity were performed as described previously [10].

### Behavioral sleep analysis

Behavioral data for flies in imaging experiments was analyzed as previously [10] using a custom-written MatLab code that measured the pixel change occurring over the legs of the fly on the ball over the entire time series. Data was analyzed and graphed using Graphpad Prism. All data was checked for Gaussian distribution using a D’Agostino-Pearson normality test prior to statistical testing. Data from THIP experiments was analyzed using a non-parametric Mann-Whitney test.

Sleep behavior in freely-walking flies was analyzed with the Drosophila ARousal Tracking system (DART) as previously described [84]. Prior to analysis, 3-5 day-old females were collected and loaded individually into 65 mm glass tubes (Trikinetics) that were plugged at one end with our standard fly food (see above), containing either 0.1 mg/ml THIP or 0.5 mg/ml all-trans-retinal (ATR). Controls were placed onto normal food and housed under identical conditions as the experimental groups. The tubes were placed onto platforms (6 total platforms, 17 tubes per platform, up to 102 flies total) for filming. THIP-fed flies were monitored for 3 days/nights, and sleep data were averaged as no significant differences were detected across successive days. Optogenetically-activated were tracked for baseline day without red light, after which they were exposed to ultra-bright red LED (617 nm Luxeon Rebel LED, Luxeon Star LEDs, Ontario, Canada) which produce 0.1- 0.2mW/mm2 at a distance of 4-5 cm with the aid of 723 concentrator optics (Polymer Optics 6° 15 mm Circular Beam Optic, Luxeon Star LEDs) for the duration of the experiment for optogenetic activation. Significance was determined by ANOVA with Tukey’s multiple comparisons test (GraphPad Prism). Sleep analysis in nAchRα knockout animals was performed using Trikinetics beam-crossing devices, with regular (>5min) and short sleep (1- 5min) calculated as previously [10].

### Sleep deprivation

Flies were sleep deprived (SD) with the use of the previously described Sleep Nullifying Apparatus (SNAP), an automated sleep deprivation apparatus that has been found to keep flies awake without nonspecifically activating stress responses [37]. Vials containing no greater than 20 flies, which contained either standard food medium or medium containing 0.1mg/ml THIP were placed on the SNAP apparatus for continuous sleep deprivation. The SNAP apparatus was programmed to snap the flies once every 20 seconds for the duration of the sleep deprivation protocol.

### RNA-Sequencing

Flies collected for RNA-sequencing analysis were first housed in vials containing either 0.5mg/ml all-trans retinal (ATR) or 0.1mg/ml THIP for sleep induction, along with their genetically identical controls on standard food medium. Flies undergoing sleep induction by optogenetic activation with ATR and their controls were placed under constant red-light from 8AM until 6PM to coincide with normal 12:12 light/dark cycles. Flies were collected after 1 hour (ZT 1) and 10 hours (ZT 10) post induction for immediate brain dissection and RNA extraction. For analysis of pharmacological sleep induction, flies were placed on THIP or normal food medium at 8AM (ZT 0) and collected for dissection at 6PM (ZT 10).

Whole fly brains were dissected in ice cold RNA*later (Sigma-Aldrich)* with 0.1% PBST as per previously published protocol [85]. The dissected brains were immediately pooled into five 1.5-mL Eppendorf tubes containing 5 brains (*n* = 25) each. Total RNA was immediately purified using TRIzol according to the manufacturer’s protocols (Sigma-Aldrich) and stored at -80°C until commencement of RNA-sequencing.

cDNA libraries were prepared using the Illumina TruSeq stranded mRNA library prep kit. Image processing and sequence data extraction were performed using the standard Illumina Genome Analyzer software and CASAVA (version 1.8.2) software. Cutadapt (version 1.8.1) was used to cut the adaptor sequences as well as low quality nucleotides at both ends. When a processed read is shorter than 36 bp, the read was discarded by cutadapt, with the parameter setting of “-q 20,20 --minimum-length=36”. Processed reads were aligned to the *Drosophila melanogaster* reference genome (dm6) using HISAT2 (version 2.0.5) [86], with the parameter setting of “--no-unal --fr --rna-strandness RF --known-splicesite-infile dm6_splicesites.txt”. This setting is to i) suppress SAM records for reads that failed to align (“--no-unal”), ii) specify the Illumina’s paired-end sequencing assay and the strand-specific information (“--fr --rna-strandness RF”) and iii) provide a list of known splice sites in *Drosophila melanogaster* (“--known-splicesite-infile dm6_splicesites.txt”). Samtools (version 1.3) [87] was then used to convert “SAM” files to “BAM” files, sort and index the “BAM” files. The “htseq-count” module in the HTSeq package (v0.7.1) was used to quantitate the gene expression level by generating a raw count table for each sample (i.e. counting reads in gene features for each sample). Based on these raw count tables, edgeR (version 3.16.5) [88] was adopted to perform the differential expression analysis between treatment groups and controls. EdgeR used a trimmed mean of M-values to compute scale factors for library size normalization [89]. It used the Cox-Reid profile-adjusted likelihood method to estimate dispersions [90] and the quasi-likelihood F-test to determine differential expression [91]. Lowly expressed genes in both groups (the mean CPM < 5 in both groups) were removed. Differentially expressed genes were identified using the following criteria: i) FDR < 0.05 and ii) fold changes > 1.5 (or logfc >0.58). Gene ontology enrichment analysis for differentially expressed genes was performed using the functional annotation tool in DAVID Bioinformatics Resources (version 6.8) [92, 93].

### Gene expression

#### RNA and cDNA Synthesis

A quantitative reverse transcriptase PCR assay was used to confirm expression of genes enriched during THIP sleep induction. Nineteen candidate genes were selected (eight negatively and eleven positively) for the gaboxadol (THIP) sleep analysis and six genes (four negatively and two positively) for the dFSB activation experiments. Total RNA was isolated using the Directzol RNA kit (ZymoResearch) from twenty adult brains per condition and each condition was collected in triplicate (i.e., 3 biological replicates). RNA quality was confirmed using a microvolume spectrophotometer NanoDrop 2000 (Thermo, USA) with only those resulting samples meeting optimal density ratios between 1.8 and 2.1 used. Up to 1 μg of total RNA was reverse transcribed using a High-Capacity cDNA Reverse Transcription Kit (Themo, USA) as per the manufacturer’s protocols. The synthesis of cDNA and subsequent amplification was performed in max volumes of 20 μL per reaction using the T100 Thermal Cycler (Bio-Rad, USA). Thermocycle conditions were as such; 25 °C for 10 min, 37 °C for 120 min, 85 °C for 5 min, and held at 4 °C. All cDNA was subsequently stored at − 20 °C until used. Target genes for THIP experiments included Pxt (CG7660, FBgn0261987), RpS5b (CG7014, FBgn0038277), Dhd (CG4193, FBgn0011761), CG9377 (CG9377, FBgn0032507), aKHr (CG11325, FBgn0025595), Acox57D-d (CG9709, FBgn0034629), FASN1 (CG3523, FBgn0283427), Pen (CG4799, FBgn0287720), CG10513 (CG10513, FBgn0039311), Gasp (CG10287, FBgn0026077), Act57B (CG10067, FBgn0000044), Bin1 (CG6046, FBgn0024491), verm (CG8756, FBgn0261341), CG16885 (CG16885, FBgn0032538), CG16884 (CG16884, FBgn0028544), CG5999 (CG5999, FBgn0038083), Fbp1 (CG17285, FBgn0000639), CG5724 (CG5724, FBgn0038082), Eh (CG5400, FBgn0000564). Target genes for dFSB experiments included Vmat (CG33528, FBgn0260964), Dop1R1 (CG9652, FBgn0011582), Salt (CG2196, FBgn0039872), Dysb (CG6856, FBgn0036819), Irk3 (CG10369, FBgn0032706), Blos1 (CG30077, FBgn0050077). Housekeeping genes included Rpl32 (CG7939, FBgn0002626), Gapdh2 (CG8893, FBgn0001092), Actin 5C (CG4027, FBgn0000042). Primer sequences can be found in **Supplementary File 3**.

#### Quantitative real-time PCR

Quantitative (q) RT-PCR was carried out using the Luna Universal qPCR Master Mix (NEB) in the CFX384 Real-Time system (Bio-Rad, USA). Cycling conditions were: 1. 95°C for 60 s, 2. 95°C for 15s, 3. 60°C for 60s with 39 cycles of steps two and three. Melt curve analysis was then performed with the following conditions 1. 95°C for 15s, 2. 60°C for 60s, 3. 95°C for 15s. Three biological replicates for each condition as well as three technical replicates per biological sample were loaded. Each experiment was then repeated on three separate occasions. Cq values and standard curves were generated using Bio Rad CFX Manager Software to ensure amplification specificity. Results were normalized to the above housekeeping genes and gene expression was calculated following the 2^− ΔΔCq method (Livak and Schmittgen 2001). All primer sequenced used for these validation experiments are listed in **Supplementary File 3.**

### Gene knockouts and knockdowns

Dα1KO harboured an ends-out mediated deletion of Dα1 in a w^1118^ background with the X chromosome replaced with one from the wild type line DGRP line 59 [59]. For Dα2KO, 7kb of the Da2 locus was removed using the endogenously expressing Actin-Cas9 strain (BDSC 5490) crossed to a strain expressing two Dα2 gene locus specific sgRNAs (Dα_sgRNA_start_1 5’CATGTTTAGCGCTGCAATGC, Dα2_sgRNA_end_1 5’TTACAAGCCATCTGCCTAG). Transgenic flies were generated using the pCFD4- U6:1_U6:3tandem gRNAs construct. This was a gift from Simon Bullock. For Dα3KO (*Dα3Δ1020*), Dα4KO (*Dα4ΔBA*), Dα6KO, Dα7KO (*Dα7ΔD6*), two sgRNAs were designed to target the start and the end of the coding sequence and cloned into either pU6-BbsI-gRNA or pCFD4 plasmids. These plasmids were then microinjected into *Drosophila* embryos to generate transgenic strains stably expressing sgRNAs. These strains were crossed to another strain expressing Cas9 under Actin promoter (ActinCas9). Their offspring were screened for deletion events with PCR and crossed to appropriate balancer strains to isolate and generate homozygous knockout strains. Full deletions were identified for all these subunit genes except for Dα3 which has two partial deletions at the 3’ and 5’ ends; these were verified to be true knockouts by the lack of RNA expression [49]. ActinCas9 strain was used as genetic control for Dα3KO and Dα7KO, while this same strain with the X chromosome replaced with one from w^1118^ (w^1118^ActinCas9) was used as genetic control for Dα2KO, Dα4KO, and Dα6KO. The RNAi strain for the adipokinetic hormone receptor gene knockdown experiments (UAS-*AkhR-*RNAi) was obtained from the VDRC (KK109300). This RNAi construct was expressed in neurons across the fly brain by crossing the UAS flies to R57C10- Gal4 [30]. RNAi strains for the nicotinic alpha receptor gene knockdown experiments (UAS- nAchRα1-7-RNAi) were obtained from the BDRC. These were: nAChRα1 RNAi – BL# 28688 - y[1] v[1]; P{y[+t7.7] v[+t1.8]=TRiP.JF03103}attP2, nAChRα2 RNAi – BL# 27493 - y[1] v[1]; P{y[+t7.7] v[+t1.8]=TRiP.JF02643}attP2, nAChRα3 RNAi – BL# 27671 – y[1] v[1]; P{y[+t7.7] v[+t1.8]=TRiP.JF02750}attP2, nAChRα4 RNAi – BL# 31985 - y[1] v[1]; P{y[+t7.7] v[+t1.8]=TRiP.JF03419}attP2, nAChRα5 RNAi – BL# 77418 - y[1] sc[*] v[1] sev[21]; P{y[+t7.7] v[+t1.8]=TRiP.HMC06550}attP2, nAChRα6 RNAi – BL# 52885 - y[1] sc[*] v[1] sev[21]; P{y[+t7.7] v[+t1.8]=TRiP.HMC03623}attP40, nAChRα7 RNAi – BL# 27251 - y[1] v[1]; P{y[+t7.7] v[+t1.8]=TRiP.JF02570}attP2. These RNAi constructs were expressed in the sleep-promoting neurons by crossing the UAS flies to R23E10-Gal4.

### Code, data, and materials availability

The data and analysis tools underpinning this study (brain imaging as well as RNA sequencing) are available online via Dryad: https://doi.org/10.5061/dryad.1rn8pk10x. See **Supplemental Files 4&5** for details on accessing these datasets and code. Datasets in this study are also available from the Lead Contact upon request. All Matlab analysis code for brain imaging is the same as in a previous publication [10], and is also available from the above link or the Lead Contact. Genetic reagents used in this study are available from the Lead Contact upon request.

## Acknowledgements

This work was supported by National Health and Medical Research Council grant GNT1164499 to BvS and NIH R01 grant NS076980 to PJS and BvS.

## Supplementary Files

**Supplementary File 1 (associated with Figure 2): A comparison of sleep duration profiles (min/hr) during optogenetic and THIP induced sleep.** Tested with 2way ANOVA with Tukey’s multiple comparison test.

**Supplementary File 2 (associated with Figures 6&7).** Raw data and statistics for RT qPCR experiments.

**Supplementary File 3 (associated with Figures 6 & 7).** Primer list for RT qPCR validation experiments.

**Supplementary File 4 (associated with Materials and Methods).** Materials Design Analysis Reporting (MDAR) Checklist for authors.

**Supplementary File 5 (associated with Materials and Methods).** Readme file explaining how to access various components in the datasets made available.

## Figure Legends

**Figure 6-figure supplement 1. Gene Ontology (GO) enrichment analysis for THIP- induced sleep.** Significantly downregulated and upregulated GO categories for THIP-sleep (Figure 6-source data 1), listed from most enriched at the top. Broad GO categories are identified below.

**Figure 6-figure supplement 2. Gene Ontology (GO) enrichment analysis for THIP- provisioned flies that were sleep deprived.** Significantly downregulated and upregulated GO categories for sleep deprived flies (Figure 6-source data 2), listed from most enriched at the top. Broad GO categories are identified below.

**Figure 6-source data 1.** List of significant THIP-sleep genes. Raw data for all genes.

**Figure 6-source data 2.** List of significant sleep-deprivation genes in THIP-fed flies. Raw data for all genes.

**Figure 7-figure supplement 1. Gene Ontology (GO) enrichment analysis for optogenetic-induced sleep.** Significantly downregulated and upregulated GO categories for optogenetic-sleep (Figure 7-source data 1), listed from most enriched at the top. Broad GO categories are identified below.

**Figure 7-figure supplement 2. Circadian-related genes uncovered in optogenetic-sleep dataset. A.** Zeitgeber (ZT) 10 timepoint was compared with ZT in to uncover potential circadian-regulated genes, in two separate datasets (-ATR and +ATR). 98 genes were shared between these datasets. **B.** Of the 98 shared genes, circadian-related processes were highly enriched. **C.** Expression levels of 7 circadian genes drawn from the two different datasets in A.

**Figure 7-figure supplement 3. Summary of different Gene Ontogeny pathways engaged by optogenetic-induced sleep and THIP-induced sleep. A.** Either sleep induction method produces different levels of activity in the fly brain. We term optogenetic-induced sleep ‘active sleep’ because brain activity levels are not different than during wake. We term THIP- induced sleep ‘quiet sleep’ because neural activity is significant decreased already in the first 5 minutes. Both of these induced forms of sleep resemble sleep stages seen during spontaneous sleep in flies. **B.** Number of GO pathways engaged by either induced active or quiet sleep, separated by upregulated versus downregulated biological processes.

**Figure 7-source data 1.** List of significant optogenetic-sleep genes after 10 hours activation. Raw data for all genes.

**Figure 7-source data 2.** List of significant optogenetic-sleep genes after 1 hour of activation. Raw data for all genes.

**Figure 7-source data 3.** List of significant ZT10 vs ZT1 genes in ATR+ dataset. Raw data for all genes.

**Figure 7-source data 4.** List of significant ZT10 vs ZT1 genes in ATR- dataset. Raw data for all genes.

**Figure 8-figure supplement 1. Waking activity levels of nAchRα knockout mutants.** Top: mutants are compared to their genetic background strain. Da1-7 = nAchRα1-7. Bottom: statistical tests for waking activity levels in each knockout compared to its genetic control, during both day and night.

## References

1. Jaggard, J.B., G.X. Wang, and P. Mourrain, Non-REM and REM/paradoxical sleep dynamics across phylogeny. Curr Opin Neurobiol, 2021. 71: p. 44–51.

2. Van De Poll, M.N. and B. van Swinderen, Balancing Prediction and Surprise: A Role for Active Sleep at the Dawn of Consciousness? Front Syst Neurosci, 2021. 15: p. 768762.

3. Dijk, D.J., D.P. Brunner, and A.A. Borbely, Time course of EEG power density during long sleep in humans. Am J Physiol, 1990. 258(3 Pt 2): p. R650–61.

4. Tononi, G. and C. Cirelli, Sleep and the price of plasticity: from synaptic and cellular homeostasis to memory consolidation and integration. Neuron, 2014. 81(1): p. 12–34.

5. Siegel, J.M., REM sleep: a biological and psychological paradox. Sleep Med Rev, 2011. 15(3): p. 139–42.

6. Shein-Idelson, M., et al., Slow waves, sharp waves, ripples, and REM in sleeping dragons. Science, 2016. 352(6285): p. 590–5.

7. Rattenborg, N.C., et al., Local Aspects of Avian Non-REM and REM Sleep. Front Neurosci, 2019. 13: p. 567.

8. Libourel, P.A. and A. Herrel, Sleep in amphibians and reptiles: a review and a preliminary analysis of evolutionary patterns. Biol Rev Camb Philos Soc, 2016. 91(3): p. 833–66.

9. Yap, M.H.W., et al., Oscillatory brain activity in spontaneous and induced sleep stages in flies. Nat Commun, 2017. 8(1): p. 1815.

10. Tainton-Heap, L.A.L., et al., A Paradoxical Kind of Sleep in Drosophila melanogaster. Curr Biol, 2021. 31(3): p. 578–590 e6.

11. Leung, L.C., et al., Neural signatures of sleep in zebrafish. Nature, 2019. 571(7764): p. 198–204.

12. Iglesias, T.L., et al., Cyclic nature of the REM sleep-like state in the cuttlefish Sepia officinalis. J Exp Biol, 2019. 222(Pt 1).

13. Sauer, S., et al., The dynamics of sleep-like behaviour in honey bees. J Comp Physiol A Neuroethol Sens Neural Behav Physiol, 2003. 189(8): p. 599–607.

14. Calcagno, B., et al., Transient activation of dopaminergic neurons during development modulates visual responsiveness, locomotion and brain activity in a dopamine ontogeny model of schizophrenia. Transl Psychiatry, 2013. 2: p. e2026.

15. Shaw, P.J., et al., Correlates of sleep and waking in Drosophila melanogaster. Science, 2000. 287(5459): p. 1834–7.

16. Hendricks, J.C., et al., Rest in Drosophila is a sleep-like state. Neuron, 2000. 25(1): p. 129–38.

17. Shafer, O.T. and A.C. Keene, The Regulation of Drosophila Sleep. Curr Biol, 2021. 31(1): p. R38–R49.

18. Wolff, T. and G.M. Rubin, Neuroarchitecture of the Drosophila central complex: A catalog of nodulus and asymmetrical body neurons and a revision of the protocerebral bridge catalog. J Comp Neurol, 2018. 526(16): p. 2585–2611.

19. Troup, M., et al., Acute control of the sleep switch in Drosophila reveals a role for gap junctions in regulating behavioral responsiveness. Elife, 2018. 7.

20. Donlea, J.M., et al., Inducing sleep by remote control facilitates memory consolidation in Drosophila. Science, 2011. 332(6037): p. 1571-6.

21. Donlea, J.M., D. Pimentel, and G. Miesenbock, Neuronal machinery of sleep homeostasis in Drosophila. Neuron, 2014. 81(4): p. 860–72.

22. Donlea, J.M., et al., Recurrent Circuitry for Balancing Sleep Need and Sleep. Neuron, 2018. 97(2): p. 378–389 e4.

23. Dissel, S., et al., Sleep restores behavioral plasticity to Drosophila mutants. Curr Biol, 2015. 25(10): p. 1270–81.

24. Harrison, N.L., Mechanisms of sleep induction by GABA(A) receptor agonists. J Clin Psychiatry, 2007. 68 **Suppl 5**: p. 6–12.

25. Berry, J.A., et al., Sleep Facilitates Memory by Blocking Dopamine Neuron-Mediated Forgetting. Cell, 2015. 161(7): p. 1656–67.

26. Stanhope, B.A., et al., Sleep Regulates Glial Plasticity and Expression of the Engulfment Receptor Draper Following Neural Injury. Curr Biol, 2020. 30(6): p. 1092–1101 e3.

27. van Alphen, B., et al., A deep sleep stage in Drosophila with a functional role in waste clearance. Sci Adv, 2021. 7(4).

28. Lundahl, J., et al., EEG spectral power density profiles during NREM sleep for gaboxadol and zolpidem in patients with primary insomnia. J Psychopharmacol, 2012. 26(8): p. 1081–7.

29. Mitler, M.M. and W.C. Dement, Cataplectic-like behavior in cats after micro-injections of carbachol in pontine reticular formation. Brain Res, 1974. 68(2): p. 335–43.

30. Jenett, A., et al., A GAL4-driver line resource for Drosophila neurobiology. Cell Rep, 2012. 2(4): p. 991–1001.

31. Klapoetke, N.C., et al., Independent optical excitation of distinct neural populations. Nat Methods, 2014. 11(3): p. 338–46.

32. Zimmerman, J.E., et al., A video method to study Drosophila sleep. Sleep, 2008. 31(11): p. 1587–98.

33. Huber, R., et al., Sleep homeostasis in Drosophila melanogaster. Sleep, 2004. 27(4): p. 628–39.

34. Andretic, R. and P.J. Shaw, Essentials of sleep recordings in Drosophila: moving beyond sleep time. Methods Enzymol, 2005. 393: p. 759–72.

35. Kirszenblat, L., R. Yaun, and B. van Swinderen, Visual experience drives sleep need in Drosophila. Sleep, 2019. 42(7).

36. Troup, M., L.A.L. Tainton-Heap, and B. van Swinderen, Neural Ensemble Fragmentation in the Anesthetized Drosophila Brain. J Neurosci, 2023. 43(14): p. 2537–2551.

37. Seugnet, L., et al., Persistent short-term memory defects following sleep deprivation in a drosophila model of Parkinson disease. Sleep, 2009. 32(8): p. 984–92.

38. Muheim, C.M., et al., Dynamic- and Frequency-Specific Regulation of Sleep Oscillations by Cortical Potassium Channels. Curr Biol, 2019. 29(18): p. 2983–2992 e3.

39. He, Q., et al., AKH-FOXO pathway regulates starvation-induced sleep loss through remodeling of the small ventral lateral neuron dorsal projections. PLoS Genet, 2020. 16(10): p. e1009181.

40. Bharucha, K.N., P. Tarr, and S.L. Zipursky, A glucagon-like endocrine pathway in Drosophila modulates both lipid and carbohydrate homeostasis. J Exp Biol, 2008. 211(Pt 19): p. 3103–10.

41. Ferguson, L., et al., Transient Dysregulation of Dopamine Signaling in a Developing Drosophila Arousal Circuit Permanently Impairs Behavioral Responsiveness in Adults. Front Psychiatry, 2017. 8: p. 22.

42. Shao, L., et al., Schizophrenia susceptibility gene dysbindin regulates glutamatergic and dopaminergic functions via distinctive mechanisms in Drosophila. Proc Natl Acad Sci U S A, 2011. 108(46): p. 18831–6.

43. Van Swinderen, B. and R. Andretic, Dopamine in Drosophila: setting arousal thresholds in a miniature brain. Proc Biol Sci, 2011. 278(1707): p. 906–13.

44. Watson, C.J., H.A. Baghdoyan, and R. Lydic, >Neuropharmacology of Sleep and Wakefulness. Sleep Med Clin, 2010. 5(4): p. 513–528.

45. Rosenthal, J.S. and Q. Yuan, Constructing and Tuning Excitatory Cholinergic Synapses: The Multifaceted Functions of Nicotinic Acetylcholine Receptors in Drosophila Neural Development and Physiology. Front Cell Neurosci, 2021. 15: p. 720560.

46. Shi, M., et al., Identification of Redeye, a new sleep-regulating protein whose expression is modulated by sleep amount. Elife, 2014. 3: p. e01473.

47. Dai, X., et al., Molecular resolution of a behavioral paradox: sleep and arousal are regulated by distinct acetylcholine receptors in different neuronal types in Drosophila. Sleep, 2021. 44(7).

48. Chen, W., et al., Dual nicotinic acetylcholine receptor subunit gene knockouts reveal limits to functional redundancy. Pestic Biochem Physiol, 2022. 184: p. 105118.

49. Perry, T., et al., Role of nicotinic acetylcholine receptor subunits in the mode of action of neonicotinoid, sulfoximine and spinosyn insecticides in Drosophila melanogaster. Insect Biochem Mol Biol, 2021. 131: p. 103547.

50. van Alphen, B., et al., A dynamic deep sleep stage in Drosophila. J Neurosci, 2013. 33(16): p. 6917–27.

51. Xie, L., et al., Sleep drives metabolite clearance from the adult brain. Science, 2013. 342(6156): p. 373–7.

52. Kubin, L., Carbachol models of REM sleep: recent developments and new directions. Arch Ital Biol, 2001. 139(1-2): p. 147–68.

53. Torterolo, P., et al., Neocortical 40 Hz oscillations during carbachol-induced rapid eye movement sleep and cataplexy. Eur J Neurosci, 2016. 43(4): p. 580–9.

54. Kirszenblat, L. and B. van Swinderen, The Yin and Yang of Sleep and Attention. Trends Neurosci, 2015. 38(12): p. 776–86.

55. Vazquez, J. and H.A. Baghdoyan, Basal forebrain acetylcholine release during REM sleep is significantly greater than during waking. Am J Physiol Regul Integr Comp Physiol, 2001. 280(2): p. R598–601.

56. Smarandache-Wellmann, C.R., *Arthropod neurons and nervous system.* Curr Biol, 2016. **26**(20): p. R960–R965.

57. Bourgin, P., et al., Induction of rapid eye movement sleep by carbachol infusion into the pontine reticular formation in the rat. Neuroreport, 1995. 6(3): p. 532–6.

58. Kashiwagi, M. and Y. Hayashi, Life Without Dreams: Muscarinic Receptors Are Required to Regulate REM Sleep in Mice. Cell Rep, 2018. 24(9): p. 2211–2212.

59. Somers, J., et al., Pleiotropic Effects of Loss of the Dalpha1 Subunit in Drosophila melanogaster: Implications for Insecticide Resistance. Genetics, 2017. 205(1): p. 263–271.

60. Jones, J.D., et al., Regulation of sleep by cholinergic neurons located outside the central brain in Drosophila. PLoS Biol, 2023. 21(3): p. e3002012.

61. Wiggin, T.D., et al., Covert sleep-related biological processes are revealed by probabilistic analysis in Drosophila. Proc Natl Acad Sci U S A, 2020. 117(18): p. 10024–10034.

62. Gong, N.N., et al., Intrinsic maturation of sleep output neurons regulates sleep ontogeny in Drosophila. Curr Biol, 2022. 32(18): p. 4025–4039 e3.

63. Stahl, B.A., et al., Sleep-Dependent Modulation of Metabolic Rate in Drosophila. Sleep, 2017. 40(8).

64. Kempf, A., et al., A potassium channel beta-subunit couples mitochondrial electron transport to sleep. Nature, 2019. 568(7751): p. 230–234.

65. Liu, S., et al., Sleep Drive Is Encoded by Neural Plastic Changes in a Dedicated Circuit. Cell, 2016. 165(6): p. 1347–1360.

66. Lei, Z., K. Henderson, and K. Keleman, A neural circuit linking learning and sleep in Drosophila long-term memory. Nat Commun, 2022. 13(1): p. 609.

67. Zimmerman, J.E., et al., Conservation of sleep: insights from non-mammalian model systems. Trends Neurosci, 2008. 31(7): p. 371–6.

68. Raizen, D.M., et al., Lethargus is a Caenorhabditis elegans sleep-like state. Nature, 2008. 451(7178): p. 569–72.

69. Hill, A.J., et al., Cellular stress induces a protective sleep-like state in C. elegans. Curr Biol, 2014. 24(20): p. 2399–405.

70. Artiushin, G. and A. Sehgal, The Glial Perspective on Sleep and Circadian Rhythms. Annu Rev Neurosci, 2020. 43: p. 119–140.

71. Caron, A.M. and R. Stephenson, Energy expenditure is affected by rate of accumulation of sleep deficit in rats. Sleep, 2010. 33(9): p. 1226–35.

72. Maquet, P., Sleep function(s) and cerebral metabolism. Behav Brain Res, 1995. 69(1- 2): p. 75–83.

73. Rossler, D.C., et al., Regularly occurring bouts of retinal movements suggest an REM sleep-like state in jumping spiders. Proc Natl Acad Sci U S A, 2022. 119(33): p. e2204754119.

74. Hutchison, I.C. and S. Rathore, The role of REM sleep theta activity in emotional memory. Front Psychol, 2015. 6: p. 1439.

75. Harrington, M.O., et al., The influence of REM sleep and SWS on emotional memory consolidation in participants reporting depressive symptoms. Cortex, 2018. 99: p. 281–295.

76. Anderson, D.J. and R. Adolphs, A framework for studying emotions across species. Cell, 2014. 157(1): p. 187–200.

77. Seth, A.K. and K.J. Friston, Active interoceptive inference and the emotional brain. Philos Trans R Soc Lond B Biol Sci, 2016. 371(1708).

78. Hobson, J.A., REM sleep and dreaming: towards a theory of protoconsciousness. Nat Rev Neurosci, 2009. 10(11): p. 803–13.

79. Pribbenow, C., et al., Postsynaptic plasticity of cholinergic synapses underlies the induction and expression of appetitive and familiarity memories in Drosophila. Elife, 2022. 11.

80. Barnstedt, O., et al., Memory-Relevant Mushroom Body Output Synapses Are Cholinergic. Neuron, 2016. 89(6): p. 1237–1247.

81. Melnattur, K., et al., A conserved role for sleep in supporting Spatial Learning in Drosophila. Sleep, 2020.

82. Pfeiffer, B.D., J.W. Truman, and G.M. Rubin, Using translational enhancers to increase transgene expression in Drosophila. Proc Natl Acad Sci U S A, 2012. 109(17): p. 6626–31.

83. Legland, D., I. Arganda-Carreras, and P. Andrey, MorphoLibJ: integrated library and plugins for mathematical morphology with ImageJ. Bioinformatics, 2016. 32(22): p. 3532–3534.

84. Faville, R., et al., How deeply does your mutant sleep? Probing arousal to better understand sleep defects in Drosophila. Scientific Reports, 2015. 5: p. 8454.

85. Chen, C., et al., Drosophila Ionotropic Receptor 25a mediates circadian clock resetting by temperature. Nature, 2015. 527(7579): p. 516–20.

86. Kim, D., B. Langmead, and S.L. Salzberg, HISAT: a fast spliced aligner with low memory requirements. Nat Methods, 2015. 12(4): p. 357–60.

87. Li, H., et al., The Sequence Alignment/Map format and SAMtools. Bioinformatics, 2009. 25(16): p. 2078–9.

88. Robinson, M.D., D.J. McCarthy, and G.K. Smyth, edgeR: a Bioconductor package for differential expression analysis of digital gene expression data. Bioinformatics, 2010. 26(1): p. 139–40.

89. Robinson, M.D. and A. Oshlack, A scaling normalization method for differential expression analysis of RNA-seq data. Genome Biol, 2010. 11(3): p. R25.

90. McCarthy, D.J., Y. Chen, and G.K. Smyth, Differential expression analysis of multifactor RNA-Seq experiments with respect to biological variation. Nucleic Acids Res, 2012. 40(10): p. 4288–97.

91. Lun, A.T., Y. Chen, and G.K. Smyth, It’s DE-licious: A Recipe for Differential Expression Analyses of RNA-seq Experiments Using Quasi-Likelihood Methods in edgeR. Methods Mol Biol, 2016. 1418: p. 391–416.

92. Huang da, W., B.T. Sherman, and R.A. Lempicki, Bioinformatics enrichment tools: paths toward the comprehensive functional analysis of large gene lists. Nucleic Acids Res, 2009. 37(1): p. 1–13.

93. Huang da, W., B.T. Sherman, and R.A. Lempicki, Systematic and integrative analysis of large gene lists using DAVID bioinformatics resources. Nat Protoc, 2009. 4(1): p. 44–57.

