## Supplementary material for "Experimentally induced active and quiet sleep engage non-overlapping transcriptional programs in *Drosophila*": Figure6-figure supplement 1

Supplementary Figure 1

| | GO Term | Pvalue | Enrichment value |
|---|---|---|---|
| Downregulated | GO:0046460 neutral lipid biosynthetic process | 0.0006 | 52.92 |
| | GO:0046463 acylglycerol biosynthetic process | 0.0006 | 52.92 |
| | GO:0002181 cytoplasmic translation | 0.0000 | 33.41 |
| | GO:0009059 macromolecule biosynthetic process | 0.0000 | 6.44 |
| | GO:0009058 biosynthetic process | 0.0000 | 4.53 |
| | GO:0009064 glutamine family amino acid metabolic process | 0.0008 | 16.54 |
| | GO:0006412 translation | 0.0000 | 11.7 |
| | GO:0043043 peptide biosynthetic process | 0.0000 | 11.54 |
| | GO:0043604 amide biosynthetic process | 0.0000 | 10.48 |
| | GO:0006518 peptide metabolic process | 0.0000 | 8.89 |
| | GO:1901566 organonitrogen compound biosynthetic process | 0.0000 | 6.38 |
| | GO:0044271 cellular nitrogen compound biosynthetic process | 0.0000 | 5.07 |
| | GO:0034641 cellular nitrogen compound metabolic process | 0.0000 | 2.31 |
| | GO:1901564 organonitrogen compound metabolic process | 0.0000 | 2.06 |
| | GO:0006807 nitrogen compound metabolic process | 0.0002 | 1.6 |
| | GO:1901576 organic substance biosynthetic process | 0.0000 | 4.53 |
| | GO:0019752 carboxylic acid metabolic process | 0.0008 | 3.36 |
| | GO:0043436 oxoacid metabolic process | 0.0010 | 3.26 |
| | GO:0006082 organic acid metabolic process | 0.0010 | 3.25 |
| | GO:0019538 protein metabolic process | 0.0000 | 2.27 |
| | GO:0071704 organic substance metabolic process | 0.0000 | 1.9 |
| | GO:0043170 macromolecule metabolic process | 0.0001 | 1.72 |
| | GO:0006591 ornithine metabolic process | 0.0006 | 52.92 |
| | GO:0006525 arginine metabolic process | 0.0008 | 44.1 |
| | GO:1901605 alpha-amino acid metabolic process | 0.0003 | 6.73 |
| | GO:0008152 metabolic process | 0.0000 | 1.83 |
| | GO:0044238 primary metabolic process | 0.0000 | 1.89 |
| | GO:0019432 triglyceride biosynthetic process | 0.0003 | 66.15 |
| | GO:0006414 translational elongation | 0.0000 | 34.51 |
| | GO:0034645 cellular macromolecule biosynthetic process | 0.0000 | 7.55 |
| | GO:0043603 cellular amide metabolic process | 0.0000 | 7.7 |
| | GO:0044249 cellular biosynthetic process | 0.0000 | 4.69 |
| | GO:0044267 cellular protein metabolic process | 0.0000 | 2.58 |
| | GO:0044260 cellular macromolecule metabolic process | 0.0000 | 2.02 |
| | GO:0044237 cellular metabolic process | 0.0000 | 1.67 |
| | GO:0007548 sex differentiation | 0.0003 | 22.05 |
| Upregulated | GO:0006030 chitin metabolic process | 0.0003 | 21.44 |
| | GO:1901071 glucosamine-containing compound metabolic process | 0.0004 | 19.78 |
| | GO:0006040 amino sugar metabolic process | 0.0004 | 19.33 |
| | GO:0006022 aminoglycan metabolic process | 0.0006 | 17.48 |
| | GO:0018990 ecdysis, chitin-based cuticle | 0.0002 | 85.05 |
| | GO:0022404 molting cycle process | 0.0004 | 65.42 |
| | GO:0040003 chitin-based cuticle development | 0.0000 | 32.89 |
| | GO:0042335 cuticle development | 0.0000 | 31.33 |
| | GO:0048856 anatomical structure development | 0.0004 | 4.39 |

**Metabolic process**

- **Biosynthetic metabolic process**
- **Nitrogen compound metabolic process**
- **organic substance metabolic process**
- **primary metabolic process**
- **Cellular metabolic process**

**Developmental Process**

- **Developmental process involved in reproduction**
- **Anatomical structure development**

**Mutlicellular organismal process**

- **Molting cycle process**
