## Supplementary material for "Experimentally induced active and quiet sleep engage non-overlapping transcriptional programs in *Drosophila*": Figure6-figure supplement 2

| | GO Term | Pvalue | Enrichment value |
|---|---|---|---|
| Downregulated | GO:0104004 cellular response to environmental stimulus | 0.0005 | 19.04 |
| | GO:0050896 response to stimulus | 0.0000 | 3.79 |
| | GO:0034644 cellular response to UV | 0.0000 | 85.05 |
| | GO:0009411 response to UV | 0.0001 | 41.15 |
| | GO:0071482 cellular response to light stimulus | 0.0001 | 32.71 |
| | GO:0071478 cellular response to radiation | 0.0003 | 22.78 |
| | GO:0071214 cellular response to abiotic stimulus | 0.0005 | 19.04 |
| | GO:0009266 response to temperature stimulus | 0.0000 | 16.88 |
| | GO: 0009617 response to bacterium | 0.0000 | 21.07 |
| | GO: 0051707 response to other organism | 0.0000 | 15.39 |
| | GO:0043207 response to external biotic stimulus | 0.0000 | 15.14 |
| | GO: 0009607 response to biotic stimulus | 0.0000 | 15.09 |
| | GO:0050830 defense response to Gram-positive bacterium | 0.0000 | 36.98 |
| | GO:0009605 response to external stimulus | 0.0000 | 9.31 |
| | GO:0034605 cellular response to heat | 0.0000 | 60.75 |
| | GO:0009408 response to heat | 0.0000 | 26.58 |
| | GO:0006979 response to oxidative stress | 0.0002 | 13.83 |
| | GO:0042742 defense response to bacterium | 0.0000 | 12.89 |
| | GO:0098542 defense response to other organism | 0.0000 | 11.63 |
| | GO:0006952 defense response | 0.0000 | 10.12 |
| | GO:0006950 response to stress | 0.0000 | 5.79 |
| | GO:0042381 hemolymph coagulation | 0.0001 | 141.75 |
| | GO:0006955 immune response | 0.0007 | 9.83 |
| | GO:0002376 immune system process | 0.0002 | 9.01 |
| | GO:0007599 hemostasis | 0.0001 | 141.75 |
| | GO:0050878 regulation of body fluid levels | 0.0005 | 56.7 |
| | GO:0051704 multi-organism process | 0.0000 | 12.34 |
| | GO:0050817 coagulation | 0.0001 | 141.75 |
| Upregulated | GO:0008152 metabolic process | 0.0000 | 1.86 |
| | GO:0006032 chitin catabolic process | 0.0001 | 36.6 |
| | GO:1901072 glucosamine-containing compound catabolic process | 0.0001 | 32.53 |
| | GO:0006022 aminoglycan metabolic process | 0.0000 | 13.37 |
| | GO:0006807 nitrogen compound metabolic process | 0.0000 | 1.94 |
| | GO:0046348 amino sugar catabolic process | 0.0001 | 32.53 |
| | GO:0006026 aminoglycan catabolic process | 0.0003 | 21.69 |
| | GO:0006030 chitin metabolic process | 0.0000 | 16.4 |
| | GO:1901071 glucosamine-containing compound metabolic process | 0.0000 | 15.13 |
| | GO:0006040 amino sugar metabolic process | 0.0000 | 14.79 |
| | GO:0006508 proteolysis | 0.0000 | 5.13 |
| | GO:1901135 carbohydrate derivative metabolic process | 0.0002 | 3.94 |
| | GO:1901564 organonitrogen compound metabolic process | 0.0000 | 2.31 |
| | GO:0043170 macromolecule metabolic process | 0.0000 | 2.1 |
| | GO:0071704 organic substance metabolic process | 0.0000 | 1.92 |
| | GO:0019538 protein metabolic process | 0.0008 | 2.01 |
| | GO:0044238 primary metabolic process | 0.0004 | 1.67 |
| | GO:0017144 drug metabolic process | 0.0000 | 6.24 |

**Response to stimulus**
- **Cellular response to stimulus**
- **Response to abiotic stimulus**
- **Response to biotic stimulus**
- **Response to external stimulus**
- **Response to stress**

**Mutli-organism process**
- **Mutliorganism cellular process**

**MutliCellular organismal process**
- **Coagulation**

**Immune system process**
- **Immune response**

**Biological regulation**
- **Regulation of biological quality**

**Metabolic process**
- **Metabolic process**
- **Nitrogen compound metabolic process**
- **Organic substance**
- **Primary Metabolic process**
- **Other**
