## Supplementary material for "Experimentally induced active and quiet sleep engage non-overlapping transcriptional programs in *Drosophila*": Figure7-figure supplement 1

Supplementary Figure 3

| | GO Term | Pvalue | Enrichment value |
|---|---|---|---|
| Downregulated | GO:0060148 positive regulation of posttranscriptional gene silencing | 0.0005 | 53.51 |
| | GO:0002181 cytoplasmic translation | 0 | 9.46 |
| | GO:0052803 imidazole-containing compound metabolic process | 0.0002 | 89.19 |
| | GO:0001692 histamine metabolic process | 0.0002 | 89.19 |
| | GO:0042133 neurotransmitter metabolic process | 0.0007 | 16.72 |
| | GO:1900368 regulation of RNA interference | 0.0001 | 133.79 |
| | GO:1900370 positive regulation of RNA interference | 0.0001 | 133.79 |
| | GO:0036466 synaptic vesicle recycling via endosome | 0.0003 | 66.89 |
| | GO:0036465 synaptic vesicle recycling | 0.0008 | 44.6 |
| Upregulated | GO:2000766 negative regulation of cytoplasmic translation | 0.0003 | 63.5 |
| | GO:0017148 negative regulation of translation | 0.0002 | 9.16 |
| | GO:0034249 negative regulation of cellular amide metabolic process | 0.0004 | 7.81 |
| | GO:0006417 regulation of translation | 0.0001 | 5.69 |
| | GO:0034248 regulation of cellular amide metabolic process | 0.0001 | 4.68 |
| | GO:0010608 posttranscriptional regulation of gene expression | 0.0001 | 4.61 |
| | GO:0010468 regulation of gene expression | 0 | 2.21 |
| | GO:0031326 regulation of cellular biosynthetic process | 0.0001 | 2.04 |
| | GO:0009889 regulation of biosynthetic process | 0.0001 | 2.04 |
| | GO:0010556 regulation of macromolecule biosynthetic process | 0.0004 | 1.98 |
| | GO:0060255 regulation of macromolecule metabolic process | 0.0001 | 1.86 |
| | GO:0019222 regulation of metabolic process | 0.0001 | 1.81 |
| | GO:0051171 regulation of nitrogen compound metabolic process | 0.0005 | 1.76 |
| | GO:0080090 regulation of primary metabolic process | 0.0006 | 1.74 |
| | GO:0031323 regulation of cellular metabolic process | 0.0007 | 1.71 |
| | GO:0046011 regulation of oskar mRNA translation | 0 | 29.31 |
| | GO:1902287 semaphorin-plexin signaling pathway involved in axon guidance | 0.0001 | 95.26 |
| | GO:1902285 semaphorin-plexin signaling pathway involved in neuron projection guidance | 0.0001 | 95.26 |
| | GO:0045876 positive regulation of sister chromatid cohesion | 0.0001 | 95.26 |
| | GO:2000305 semaphorin-plexin signaling pathway involved in reg of photoreceptor cell axon guidance | 0.0001 | 95.26 |
| | GO:0048013 ephrin receptor signaling pathway | 0.0003 | 63.5 |
| | GO:0071526 semaphorin-plexin signaling pathway | 0 | 47.63 |
| | GO:0007162 negative regulation of cell adhesion | 0.0006 | 17.86 |
| | GO:0051128 regulation of cellular component organization | 0.0002 | 2.4 |
| | GO:0048522 positive regulation of cellular process | 0.0007 | 1.88 |
| | GO:0099177 regulation of trans-synaptic signaling | 0.0001 | 5.82 |
| | GO:0050804 modulation of chemical synaptic transmission | 0.0001 | 5.82 |
| | GO:0016319 mushroom body development | 0.001 | 6.52 |
| | GO:2000026 regulation of multicellular organismal development | 0.0007 | 2.57 |
| | GO:0032502 developmental process | 0.0001 | 1.81 |
| | GO:0032879 regulation of localization | 0.0006 | 2.72 |
| | GO:2000112 regulation of cellular macromolecule biosynthetic process | 0.0004 | 1.98 |
| | GO:0050794 regulation of cellular process | 0 | 1.84 |
| | GO:0048518 positive regulation of biological process | 0.0008 | 1.8 |
| | GO:0050789 regulation of biological process | 0 | 1.7 |
| | GO:0065007 biological regulation | 0 | 1.63 |
| | GO:0042391 regulation of membrane potential | 0.0008 | 6.9 |
| | GO:0065008 regulation of biological quality | 0.0002 | 2.12 |
| | GO:0120187 positive regulation of protein localization to chromatin | 0.0001 | 95.26 |
| | GO:0008049 male courtship behavior | 0.0003 | 8.36 |
| | GO:0060179 male mating behavior | 0.0005 | 7.56 |

**Regulation of biological process**
- **Regulation of metabolic process**
- **Regulation of macromolecular metabolic process**
- **Regulation of cellular process**
- **Regulation of Signalling**
- **Regulation of development process**
- **Regulation of localisation**

**Metabolic process**
- **Biosynthetic process**
- **Nitrogen compound metabolic process**
- **cellular metabolic process**

**Biological Regulation**
- **Regulation of biological process**
- **Regulation of biological quality**

**Localisation**
- **Establishment of localisation**
- **Regulation of protein localisation**

**Behaviour**
- **Reproductive behaviour**
