## Supplementary figures and images for "Experimentally induced active and quiet sleep engage non-overlapping transcriptional programs in *Drosophila*"

### Figure7-figure supplement 2

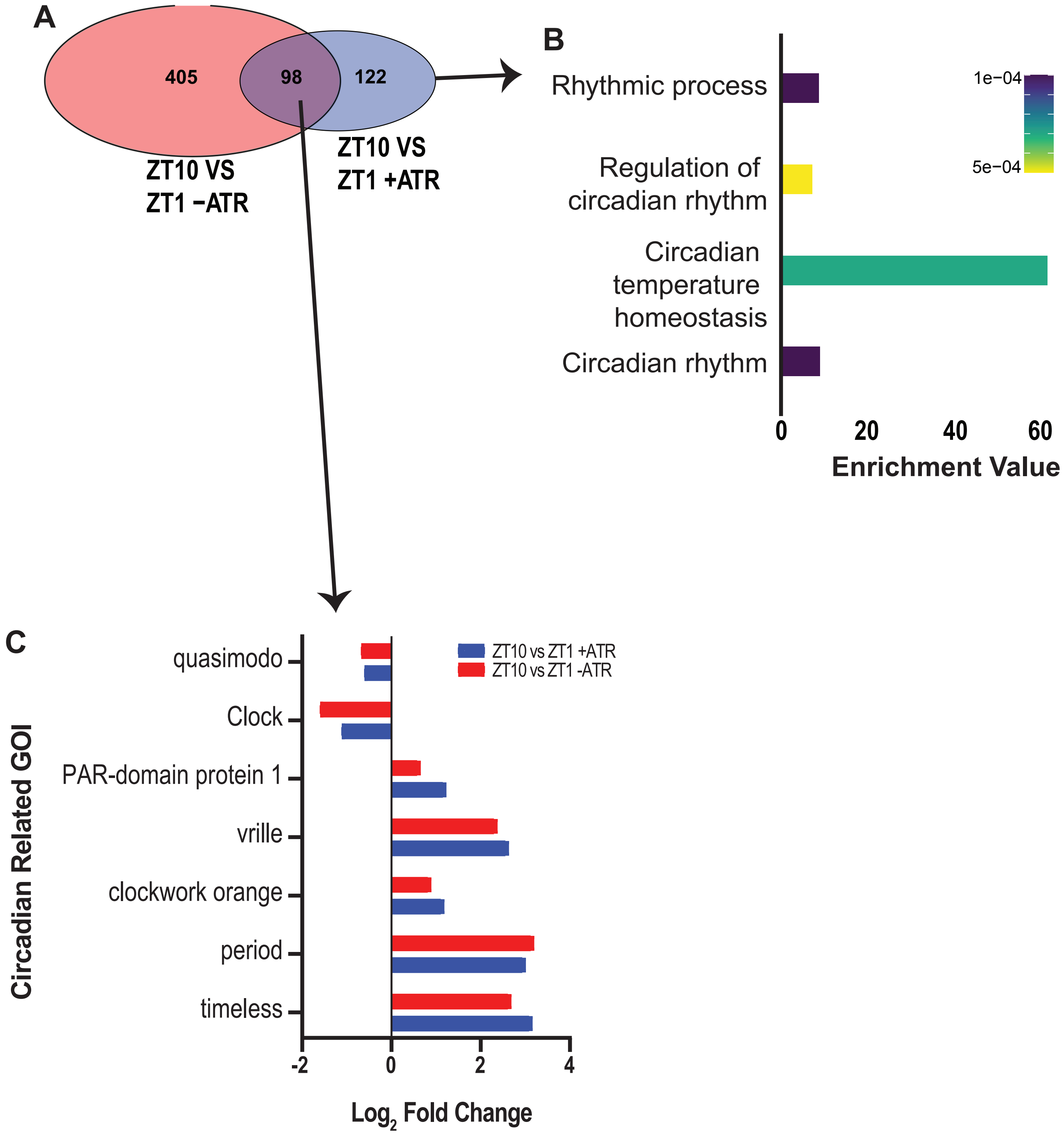
