## Supplementary material for "Experimentally induced active and quiet sleep engage non-overlapping transcriptional programs in *Drosophila*": Figure7-figure supplement 3

Supplementary Figure 5

### A: Sleep induction method

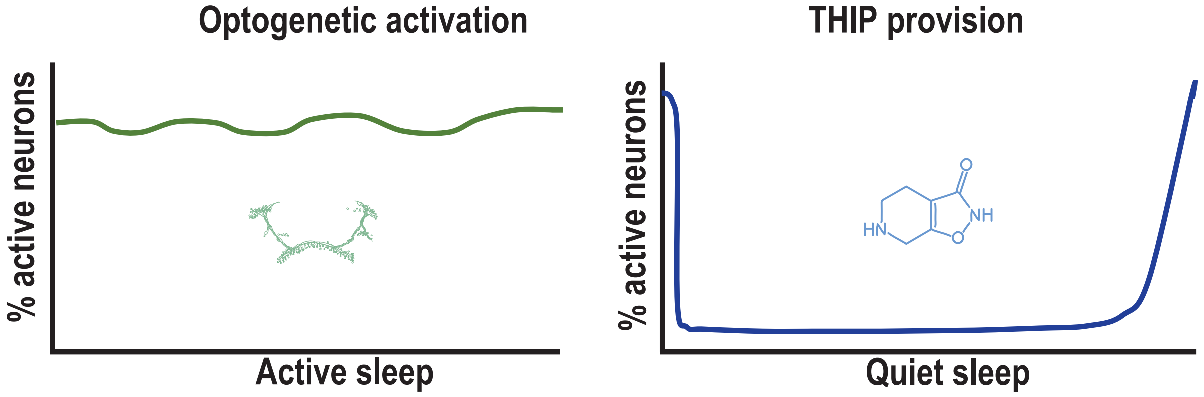


### B: Induced Sleep Transcriptome: GO Pathways of Biological Processes

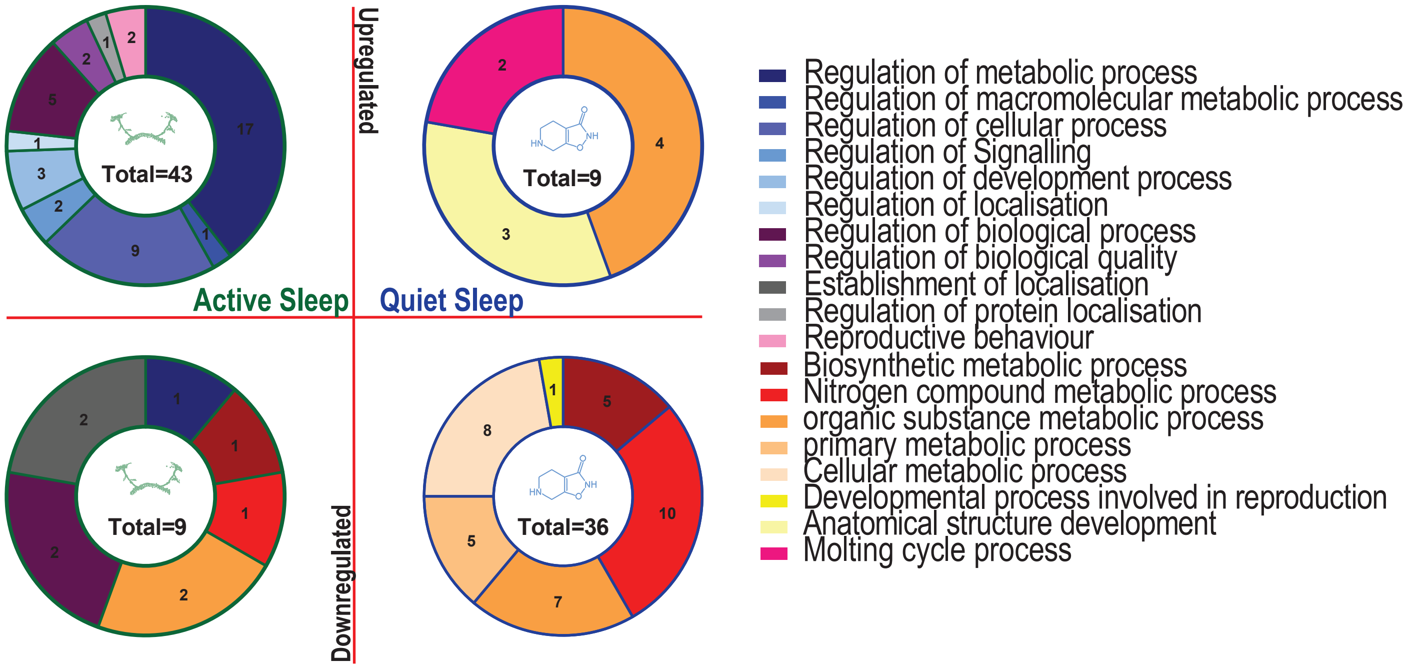
