## Supplementary material for "Experimentally induced active and quiet sleep engage non-overlapping transcriptional programs in *Drosophila*": Figure8-figure supplement 1

### Supplementary Figure 6

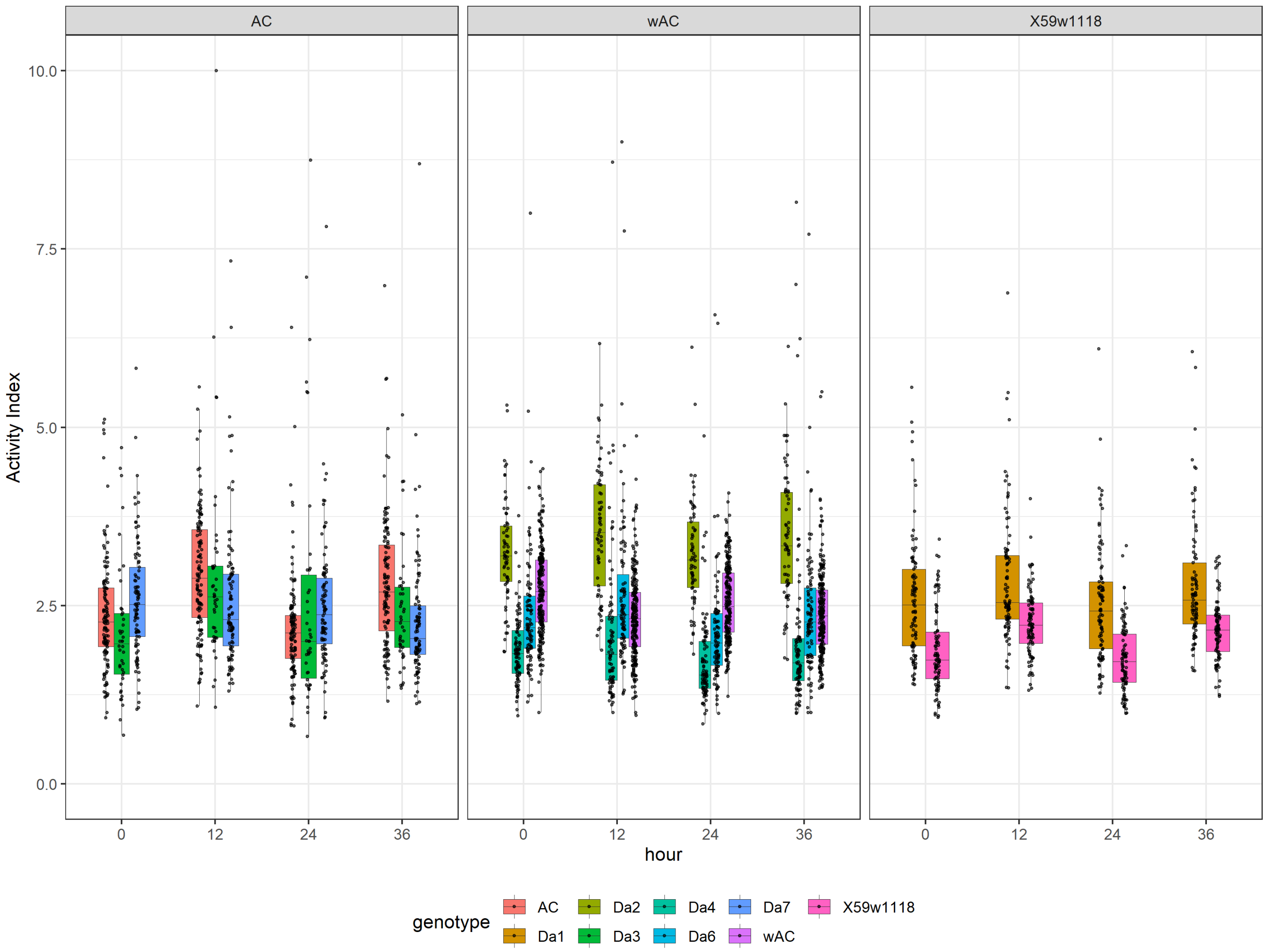


ttest_AI

| hour | gene | estimate | estimate1 | estimate2 | .y. | group1 | group2 | n1 | n2 | statistic | p | df | conf.low | conf.high | method | alternative | p.adj | p.adj.signif |
|---|---|---|---|---|---|---|---|---|---|---|---|---|---|---|---|---|---|---|
| Day | Da1 | 0.747011947979923 | 1.81085006902361 | 2.55786201700353 | activity_index | control | knockout | 91 | 93 | -7.68674248159984 | 1.63e-12 | 154.732513840803 | 0.938986358845125 | 0.555037537114721 | T-test | two.sided | 1.63e-11 | **** |
| Night | Da1 | 0.621402556761431 | 2.20459173404308 | 2.82599429080452 | activity_index | control | knockout | 91 | 93 | -6.52394430852639 | 1.39e-09 | 129.928135878636 | 0.809843317756889 | 0.432961795765974 | T-test | two.sided | 9.73e-09 | **** |
| Day | Da2 | 0.802741622150127 | 2.43199942296601 | 3.23474104511614 | activity_index | control | knockout | 70 | 65 | -7.4427805261935 | 2.03e-11 | 113.734263925721 | 1.01640704197762 | 0.589076202322632 | T-test | two.sided | 1.827e-10 | **** |
| Night | Da2 | 1.23901539806697 | 2.26258176444676 | 3.50159716251373 | activity_index | control | knockout | 70 | 65 | -10.1194970663198 | 8.52e-17 | 95.6867888875377 | 1.48206392653957 | 0.995966869594365 | T-test | two.sided | 1.1076e-15 | **** |
| Day | Da3 | 0.415534948767109 | 2.16037312206754 | 2.57590807083465 | activity_index | control | knockout | 91 | 43 | -1.34323807941607 | 0.186 | 45.4132347049576 | 1.03844775713935 | -0.207377859605138 | T-test | two.sided | 0.744 | ns |
| Night | Da3 | -0.0227331901857819 | 2.83159939569465 | 2.80886620550887 | activity_index | control | knockout | 91 | 43 | 0.0990781987694288 | 0.921 | 55.2553143772094 | 0.43704101471215 | -0.482507395083714 | T-test | two.sided | 1 | ns |
| Day | Da4 | -0.61837129538045 | 2.42331822907521 | 1.80494693369476 | activity_index | control | knockout | 98 | 87 | 8.66914228246151 | 2.56e-15 | 178.965595681112 | -0.47761492132397 | -0.75912766943693 | T-test | two.sided | 2.816e-14 | **** |
| Night | Da4 | -0.110300614309692 | 2.26586151382992 | 2.15556089952023 | activity_index | control | knockout | 98 | 87 | 0.835733550651872 | 0.405 | 115.736113589999 | 0.151109873134961 | -0.371711101754344 | T-test | two.sided | 1 | ns |
| Day | Da6 | -0.623668327284424 | 2.88534347323253 | 2.26167514594811 | activity_index | control | knockout | 91 | 91 | 6.64405236041315 | 5.29e-10 | 150.02407366578 | -0.43819297313673 | -0.809143681432117 | T-test | two.sided | 4.232e-09 | **** |
| Night | Da6 | 0.0488610995058534 | 2.54149490606688 | 2.59035600557274 | activity_index | control | knockout | 91 | 91 | -0.47619750862764 | 0.635 | 153.272240702089 | 0.251567217217754 | -0.153845018206047 | T-test | two.sided | 1 | ns |
| Day | Da7 | 0.326181724510013 | 2.22741317915403 | 2.55359490366404 | activity_index | control | knockout | 95 | 94 | -3.12088657353367 | 0.00211 | 176.4794317341 | 0.532443223747717 | 0.119920225272308 | T-test | two.sided | 0.01266 | * |
| Night | Da7 | -0.236825775153197 | 2.70854101277 | 2.4717152376168 | activity_index | control | knockout | 95 | 94 | 1.93009604618506 | 0.0552 | 181.674501771687 | 0.00527760128503382 | -0.478929151591428 | T-test | two.sided | 0.276 | ns |
