## Supplementary File 1 for "Experimentally induced active and quiet sleep engage non-overlapping transcriptional programs in *Drosophila*"

| **Tukey's multiple comparisons test** | **Mean Diff.** | **95.00% CI of diff.** | **Significant** | **Summary** | **Adjusted P Value** |
| --- | --- | --- | --- | --- | --- |
| -THIP day vs. +THIP day | -30.83 | -34.24 to -27.42 | Yes | **** | <0.0001 |
| -THIP day vs. -THIP night | -23.51 | -26.92 to -20.10 | Yes | **** | <0.0001 |
| -THIP day vs. +THIP night | -34.49 | -37.90 to -31.08 | Yes | **** | <0.0001 |
| -THIP day vs. Baseline day | -0.5857 | -3.995 to 2.824 | No | ns | 0.9995 |
| -THIP day vs. ATR day | -38.19 | -41.60 to -34.78 | Yes | **** | <0.0001 |
| -THIP day vs. Baseline night | -18.05 | -21.46 to -14.64 | Yes | **** | <0.0001 |
| -THIP day vs. ATR night | -33.84 | -37.25 to -30.43 | Yes | **** | <0.0001 |
| +THIP day vs. -THIP night | 7.315 | 3.905 to 10.72 | Yes | **** | <0.0001 |
| +THIP day vs. +THIP night | -3.665 | -7.074 to -0.2555 | Yes | * | 0.0250 |
| +THIP day vs. Baseline day | 30.24 | 26.83 to 33.65 | Yes | **** | <0.0001 |
| +THIP day vs. ATR day | -7.362 | -10.77 to -3.952 | Yes | **** | <0.0001 |
| +THIP day vs. Baseline night | 12.78 | 9.369 to 16.19 | Yes | **** | <0.0001 |
| +THIP day vs. ATR night | -3.007 | -6.417 to 0.4020 | No | ns | 0.1298 |
| -THIP night vs. +THIP night | -10.98 | -14.39 to -7.570 | Yes | **** | <0.0001 |
| -THIP night vs. Baseline day | 22.93 | 19.52 to 26.34 | Yes | **** | <0.0001 |
| -THIP night vs. ATR day | -14.68 | -18.09 to -11.27 | Yes | **** | <0.0001 |
| -THIP night vs. Baseline night | 5.463 | 2.054 to 8.872 | Yes | **** | <0.0001 |
| -THIP night vs. ATR night | -10.32 | -13.73 to -6.913 | Yes | **** | <0.0001 |
| +THIP night vs. Baseline day | 33.91 | 30.50 to 37.32 | Yes | **** | <0.0001 |
| +THIP night vs. ATR day | -3.697 | -7.106 to -0.2873 | Yes | * | 0.0229 |
| +THIP night vs. Baseline night | 16.44 | 13.03 to 19.85 | Yes | **** | <0.0001 |
| +THIP night vs. ATR night | 0.6575 | -2.752 to 4.067 | No | ns | 0.9990 |
| Baseline day vs. ATR day | -37.60 | -41.01 to -34.20 | Yes | **** | <0.0001 |
| Baseline day vs. Baseline night | -17.47 | -20.87 to -14.06 | Yes | **** | <0.0001 |
| Baseline day vs. ATR night | -33.25 | -36.66 to -29.84 | Yes | **** | <0.0001 |
| ATR day vs. Baseline night | 20.14 | 16.73 to 23.55 | Yes | **** | <0.0001 |
| ATR day vs. ATR night | 4.354 | 0.9448 to 7.764 | Yes | ** | 0.0029 |

**Table S1, related to Figure 2. A comparison of sleep duration profiles (min/hr) during optogenetic and THIP induced sleep.** Tested with 2way ANOVA with Tukey’s multiple comparison test.
