## Supplementary File 5 for "Experimentally induced active and quiet sleep engage non-overlapping transcriptional programs in *Drosophila*"

**README for datasets available at:** [**https://doi.org/10.5061/dryad.1rn8pk10x**](https://doi.org/10.5061/dryad.1rn8pk10x)

**Paper:** Experimentally induced active and quiet sleep engage non-overlapping transcriptional programs in *Drosophila*

**Authors:** Niki AnthoneyLucy A.L. Tainton-HeapHang LuongEleni NotarasAmber B. KewinQiongyi ZhaoTrent PerryPhilip BatterhamPaul J. ShawBruno van Swinderen

**This README file describes the data package accompanying the above publication.**

**Files:**

**2 Photon Data**

Files named: 2photon_FlyX_23E10Chrimson contain the raw 2photon data and .avi behavioural data for each fly. There are 5 files for each of the 9 flies. These files are affixed with:

 _Experiment

_ChanB_Preview

_Image_0001_0001

_ROIMask

_ROIs

These files relate to Figure 3C of the above publication.

Files named: 2photon_flyX_THIP contain the raw 2photon data and .avi behavioural data for each fly. There are 5 files for each of the 6 flies. These files are affixed with:

_Experiment

_ChanB_Preview

_Image_0001_0001

_ROIMask

_ROIs

These files relate to Figure 3D and 4D-4H of the above publication.

**THIP responsiveness**

Files named: THIP_responsiveness_FlyX contains the behavioural recordings of flies during behavioural responsiveness experiments. There are 7 files in total, relating to 6 flies. Fly 3 was filed in 2 parts.

These files relate to Figure 4C of the above publication.

**Matlab Codes**

File named: preprocessing.m contains the script used for preprocessing data two photon data. It performs filtering of ROIs based on ROI area, extraction of calcium signal over experiment and zscoring of calcium signal for analysis. Identifies which neurons are active in an experiment and creates masks of active ROIs. This file relates to two photon data found in figures 3 and 4 of the above paper.

File named: graph_theory_analysis.m contains the script which takes input from preprocessing.m. Calcium signals are temporally shuffled and correlated together 1000 times and the 95^th^ percentile value of shuffled correlations is used to identify neurons that are significantly correlated together over an experiment. This file relates to two photon data found in figures 3 and 4 of the above paper.

File named: Figures.m was used to create Figure 4f of the above paper.

**RNA Seq Data**

File named: THIP samples contains the fastq files for both R1 and R2 reads generated by RNA Sequencing for samples that had been treated with THIP. There are 40 files in total relating to the five biological replicates for each experiment (n=4) and 2 subsequent reads for each. These files relate to figure 6 of the above paper. Files are named in the following convention: experimentX_Y_read1fastq.gz where X would be replaced by the experiment number, and Y would be replaced by the sample number eg: 1-5.

1.       Experiment 7: which corresponds to the following experimental conditions. R23E10Gal4>UASChrimson - THIP

2.       Experiment 8: which corresponds to the following experimental conditions. R23E10Gal4>UASChrimson + THIP

3.       Experiment 9: which corresponds to the following experimental conditions. R23E10Gal4>UASChrimson + THIP + Sleep deprivation

4.       Experiment 10: which corresponds to the following experimental conditions. R23E10Gal4>UASChrimson - THIP + Sleep deprivation

File named: ATR samples contains the fastq files for both R1 and R2 reads generated by RNA Sequencing for samples that had been treated with ATR. There are 40 files in total relating to the five biological replicates for each experiment (n=4) and 2 subsequent reads for each. These files relate to figure 7 of the above paper. Files are named in the following convention: experimentX_Y_read1fastq.gz where X would be replaced by the experiment number, and Y would be replaced by the sample number eg: 1-5.

1.       Experiment 13: which corresponds to the following experimental conditions. R23E10Gal4>UASChrimson + ATR (1hr)

2.       Experiment 14: which corresponds to the following experimental conditions. R23E10Gal4>UASChrimson - ATR (1hr)

3.       Experiment 15: which corresponds to the following experimental conditions. R23E10Gal4>UASChrimson + ATR (10hr)

4.       Experiment 16: which corresponds to the following experimental conditions. R23E10Gal4>UASChrimson - ATR (10hr)
